# Cell type specific transcriptomic differences in depression show similar patterns between males and females but implicate distinct cell types and genes

**DOI:** 10.1101/2022.09.23.509254

**Authors:** Malosree Maitra, Haruka Mitsuhashi, Reza Rahimian, Anjali Chawla, Jennie Yang, Laura Fiori, Maria-Antonietta Davoli, Kelly Perlman, Zahia Aouabed, Deborah C Mash, Matthew Suderman, Naguib Mechawar, Gustavo Turecki, Corina Nagy

## Abstract

Major depressive disorder (MDD) is a common, heterogenous, and potentially serious psychiatric illness. Diverse brain cell types have been implicated in MDD etiology. Significant sexual differences exist in MDD clinical presentation and outcome, and recent evidence suggests different molecular bases for male and female MDD. We evaluated over 160,000 nuclei from 71 female and male donors, leveraging new and pre-existing single-nucleus RNA-sequencing data from the dorsolateral prefrontal cortex. Cell type specific transcriptome-wide threshold-free MDD-associated gene expression patterns were similar between the sexes, but significant differentially expressed genes (DEGs) diverged. Among 7 broad cell types and 41 clusters evaluated, microglia and parvalbumin interneurons contributed the most DEGs in females, while deep layer excitatory neurons, astrocytes, and oligodendrocyte precursors were the major contributors in males. Further, the Mic1 cluster with 38% of female DEGs and the ExN10_L46 cluster with 53% of male DEGs, stood out in the meta-analysis of both sexes.

## Introduction

Major depressive disorder (MDD) is a serious and potentially debilitating mental illness affecting 200-300 million people worldwide^1^. MDD is a leading cause of disability globally^1^ and some prominent symptoms in patients with MDD include persistent low mood, decreased interest and/or pleasure, sleep and appetite disturbances, feelings of worthlessness, and suicidal thoughts^2^. A number of genetic variants have been identified which contribute to the heritability of MDD^3^ and brain transcriptomic differences^4^ are detected in this disease, but the molecular etiology of MDD is still only partially understood.

There are known dissimilarities in the epidemiology and pathophysiology of MDD between the sexes. Notably, it is twice as prevalent in women than men^5^. Symptomatology differs in that, women are more likely to have comorbid anxiety, so-called atypical depression, and recurrent episodes, while men are more likely to have comorbid substance use disorders and to die by suicide^6–8^. Sex-specific molecular profiles in MDD and corresponding animal models are often attributed to hormonal differences either during development or in adulthood, to the contributions of sex-chromosomes, or to inherent sex-differences in the monoaminergic system or the hypothalamic-pituitary-adrenal axis (HPA), among other factors^6,9^.

Recent studies in humans have attempted to address the gap in our knowledge of molecular sex-differences in depression by examining MDD-associated sex-specific brain transcriptomic differences in human patients^7,10^. Using bioinformatic and meta-analysis approaches, combined with validation in animal models, these studies found that, overall, MDD-associated differences in brain transcriptomics are primarily sex-specific across brain regions, with very little overlap of differentially expressed genes (DEGs) and discordance in overall patterns of difference between the sexes.

Single-nucleus RNA-sequencing (snRNA-seq) can disentangle cell type specific transcriptomic contributions to complex neuropsychiatric conditions^11–15^, and our recent snRNA-seq results^16^ revealed disruptions in deep layer excitatory neurons and immature oligodendrocyte precursor cells (OPCs) in the pre-frontal cortex of males with MDD. Given the higher prevalence of MDD among women, the known sex-specific differences in MDD, and growing evidence that male and female MDD may be mediated by distinct brain molecular mechanisms, we conducted a study in a cohort of female individuals and applied an updated unified analysis pipeline to both the female and previously generated male cohorts. With a total of 71 individuals, 37 cases and 34 controls and over 160,000 single-nuclei profiled, our dataset represents the largest snRNA-seq study of the human brain in MDD to date. We found that the DEGs detected and the cell types with prominent differences were distinct in males and females. However, the overall patterns of MDD-associated gene expression difference within each cell type were consistent between the sexes. Whereas in males our analysis indicated a strong involvement of deep layer excitatory neurons, astrocytes, and OPCs – consistent with our previous report, in females we found a striking contribution of microglia and parvalbumin (PV) interneurons to MDD pathology.

## Results

### Profiling cells of the human dorsolateral prefrontal cortex (dlPFC)

snRNA-seq data was generated from the dlPFC for 20 female subjects with MDD and 18 neurotypical female controls (Figure 1a, schematic; Table 1, demographic and sample characteristics; Supplementary Table 1, sequencing metrics) and combined with previously generated data from males^16^. After pre-processing with a unified pipeline (methods: Sequencing, alignment, and generation of count matrices), we retained 160,711 high-quality nuclei with comparable contributions of sex (51% from females) and disease status (58% MDD). We used Harmony^17^ to correct for covariates, including batch effects (Supplementary Figure 1a-d), and applied the scclusteval^18^ workflow to optimize the Seurat clustering parameters (Supplementary Figure 2a-b) resulting in the identification of 41 nuclei clusters. Clusters mostly did not appear to be driven by batch, sex, or subject (Supplementary Figure 3a-e).

**Figure 1:**
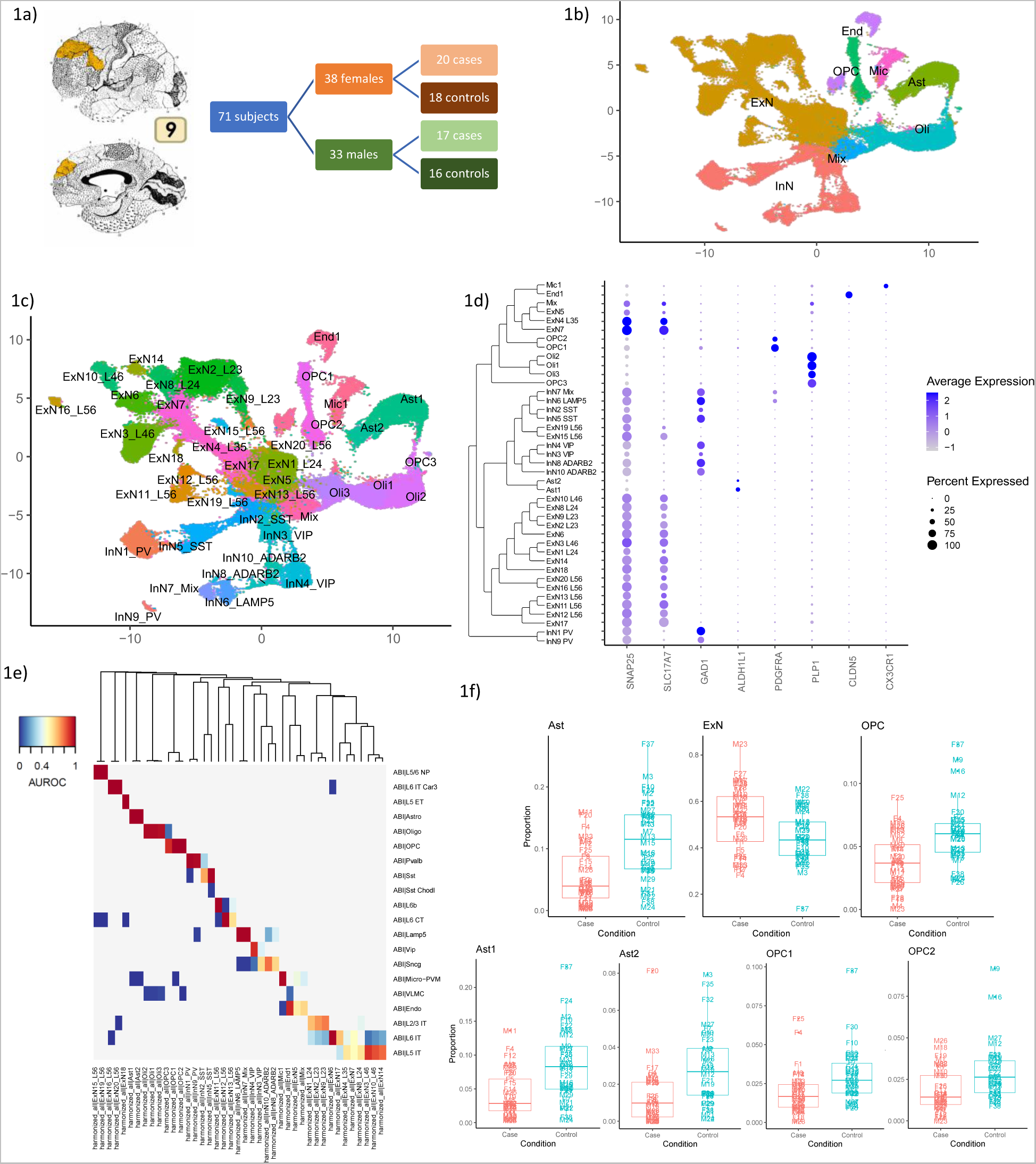
a) Schematic of study design. Diagrams depict the brain region of interest, Brodmann area 9, corresponding to the dorsolateral prefrontal cortex. b) UMAP plot colored by the broad cell types. c) UMAP plot colored by the individual clusters identified and annotated. For UMAP plots, the x and y-axes represent the first and second UMAP co-ordinates respectively. d) DotPlot depicting the expression of marker genes (*SNAP25* – neurons, *SLC17A7* – excitatory neurons, *GAD1* – inhibitory neurons, *ALDH1L1* – astrocytes, *PDGFRA* – oligodendrocyte precursor cells, *PLP1* – oligodendrocytes, *CLDN5* – endothelial cells, *CX3CR1* – microglia). The tree next to the cluster names shows the relationship between the clusters by using the distance based on average expression of highly variable genes. e) Best hits heatmap from MetaNeighbor showing the correspondence between the clusters in our dataset and the broad categories of cells identified in the Allen Brain Institute human motor cortex snRNA-seq dataset^20^. f) Boxplots showing the proportion of nuclei in each cluster for each subject split by cases and controls for the broad OPC, astrocyte, and excitatory neuron cell types and the Ast1, Ast2, OPC1, and OPC2 clusters.

**Table 1:** Demographic and sample characteristics of cohorts

| Group | Case (n = 37) |  | Control (n = 34) |  |
| --- | --- | --- | --- | --- |
| Sex | Female (n = 20) | Male (n = 17) | Female (n = 18) | Male (n = 16) |
| Age | 45.10±3.19 | 41.06±4.66 | 47.89±4.45 | 38.37±4.58 |
| PMI* | 41.49±3.07 | 41.69±4.76 | 30.27±4.73 | 32.02±4.81 |
| pH | 6.58±0.08 | 6.60±0.07 | 6.34±0.08 | 6.50±0.06 |

(\*) significantly different between female cases and controls, Kruskal-Wallis test, p-value < 0.05
Numeric values in each cell represent the mean±SEM

Of the 41 clusters, 40 could each be confidently annotated to one of 7 major brain cell types (methods: Cluster annotation, Figure 1b-d, Supplementary Figure 4, Supplementary Table 2) – excitatory neurons (48% of nuclei), inhibitory neurons (18% of nuclei), oligodendrocytes (14% of nuclei), astrocytes (8% of nuclei), OPCs (5% of nuclei), endothelial cells (2.5% of nuclei), and microglia (2% of nuclei). The one unassigned cluster displayed a mixed expression profile of neuronal and glial marker genes (2% of nuclei).

We annotated 30 neuronal clusters, both excitatory (20 clusters) and inhibitory (10 clusters), using known subtype markers (Supplementary Figure 5a-b). Excitatory neuronal clusters were annotated according to their layer of origin and inhibitory neuronal clusters according to their developmental origin, where applicable. For non-neuronal cells, we identified one microglial cluster, two clusters of astrocytes, and three clusters each of oligodendrocytes and OPCs. Clusters annotated to the oligodendrocyte lineage (OL) were further characterized using pseudotime trajectory analysis (methods: Pseudotime trajectory analysis; Supplementary Figure 5c-d).

Using the gene expression patterns of our clusters and matching them to published clusters in several human brain datasets^19,20^, we found close correspondence between observed cell types (Figure 1e, Supplementary Figure 6), further emphasizing that the quality of our data, clustering, and annotation are at par with other recent snRNA-seq datasets for the human brain.

#### Cell types with altered proportions in MDD

We next examined whether proportions of nuclei in broad cell types and clusters differed between cases and controls. We observed that the proportions of nuclei per subject contributing to the broad astrocytic and OPC cell types were significantly decreased in cases compared to controls (two-sided Wilcoxon-test, FDR =3.46×10^−4^, Ast; FDR= 5.32×10^−4^, OPC; Figure 1f, Supplementary Table 3) and there were concomitant increases in excitatory neurons (FDR 0.0477). Similarly, there were reduced proportions of nuclei in both astrocytic clusters (Ast1, FDR 0.00188; Ast2, FDR 0.00291) and in two of three OPC clusters (OPC1, FDR 0.009799; OPC2, FDR 0.0168; Figure 1f, Supplementary Table 3). The robustness of these differences was supported by sub-sampling analysis (methods: Cell-type proportions comparison). These results are similar to those found in analyses of other brain disorders^11,21^, and indicate that there may be decreased proportion of astrocytic and OPC cells may be reduced in MDD.

### Global cell type specific transcriptomic changes are largely concordant between the sexes

We next asked whether there are sex-specific differences in the gene expression patterns of individual cell types. To answer this question, we performed differential gene expression analysis comparing cases and controls in broad cell types and clusters, in males and females separately. In both males and females, we observed a high proportion of common DEGs between broad and cluster level analyses. However, consistent with previous studies showing distinct brain transcriptomic changes in males and females with MDD^10,22^, few DEGs were common to both sexes (Figure 2a). To compare overall patterns of depression-associated gene expression in males and females beyond those genes passing significance thresholds, we performed rank-rank hypergeometric overlap (RRHO) analysis^23^ (methods: Comparison of male and female results). Specifically, we used RRHO to compare the orderings of the genes induced by MDD association statistics in males compared to females. These orderings were generally strongly concordant between the sexes (Figure 2b). Evidence of discordance was visible only for Oli and OPC. There was a significant overlap between males and females in genes less expressed in MDD in Ast, ExN, and InN (warm colors in top right quadrant of RRHO plots) and an overlap in genes more expressed in MDD in Mic (warm colors in bottom left quadrant of RRHO plot).

**Figure 2:**
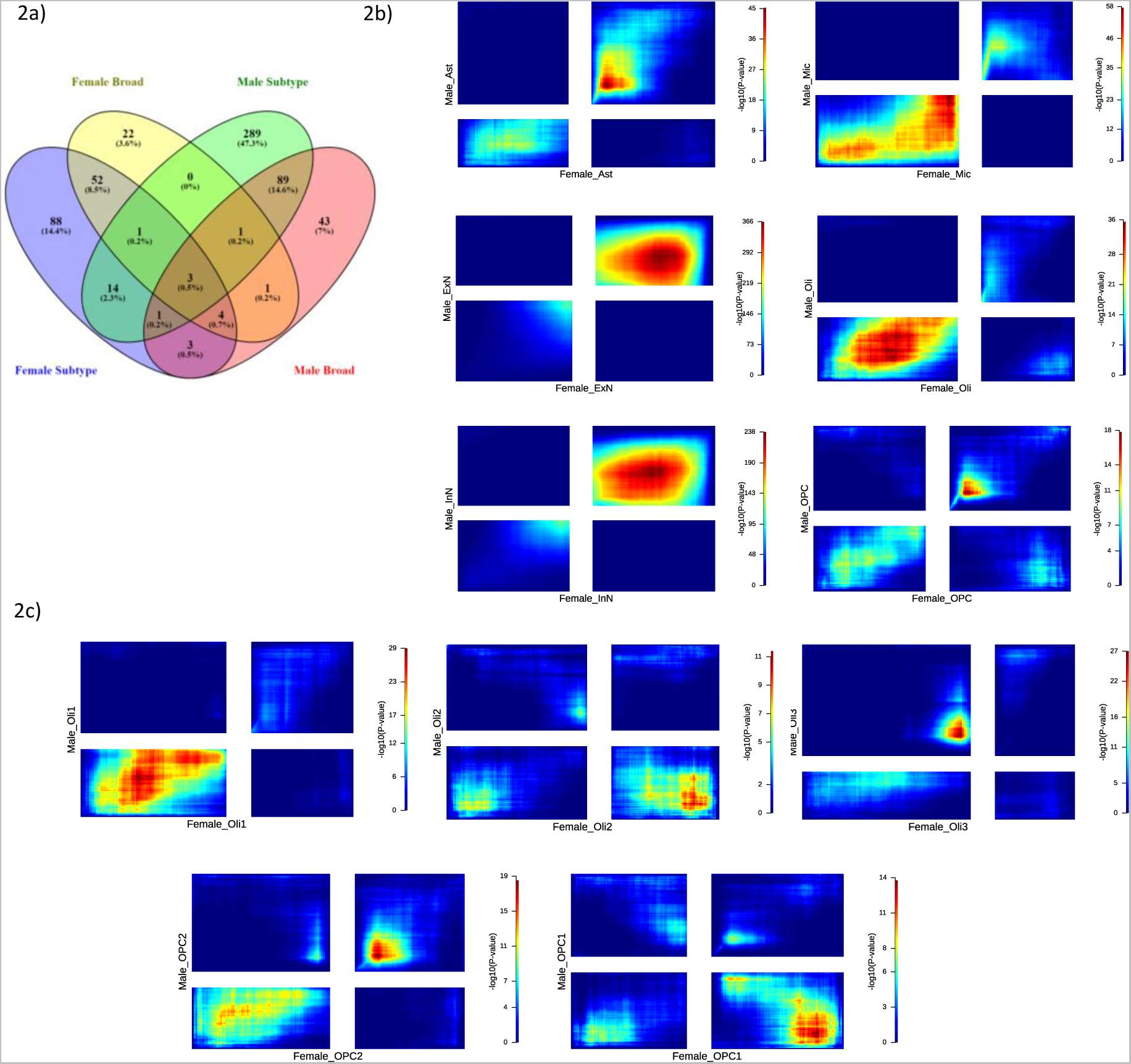
a) Venn diagram showing the overlap of DEGs between the male and female datasets at the broad cell type and cluster levels. b) RRHO2 plots for correspondence between differential expression results for broad cell types in the female (x-axis) and male (y-axis) datasets. Warm colors in the bottom left and top right quadrants reflect overlap in genes with increased expression or decreased expression respectively in cases versus controls between the male and female datasets. Warm colors in the top left and bottom right quadrants reflect overlaps in genes with opposite direction of effects between the male and female datasets. For each dataset, genes were ranked according to the value of log of fold change multiplied by the negaitve base 10 logarithm of the uncorrected p-value from differential expression analysis. c) RRHO2 plots similar to (b) but for oligodendrocyte lineage clusters.

At the cluster level there was some evidence of discordance between the sexes, with 8 out of 34 clusters compared showing discordant patterns. This encompassed certain neuronal clusters, primarily excitatory neuronal, including ExN4_L35, ExN7, ExN12_L56, ExN13_L56, InN10_ADARB2 (Supplementary Figure 7). Within the oligodendrocyte lineage, discordance is apparent for the Oli2, Oli3, and OPC1 clusters (Figure 2c).

Taken together we find that, although cell type specific statistically significant MDD-associated DEGs differ between the sexes, a threshold-free ranking approach to comparison shows considerable concordance between males and females for the majority of broad cell types and clusters.

### Cell types with strongest MDD associations differ by sex

Next, we identified the cell types with the strongest evidence of dysregulation due to MDD in each sex. In males, our reanalysis indicated results consistent with those we reported previously, i.e., for broad cell types we identified the highest number of DEGs in astrocytes (90/151, 60%) and OPCs (54/151, 36%) (Figure 3a, Supplementary Figure 8a-d, Supplementary Table 4), whereas for at the cluster level (Figure 3b, Supplementary Figure 8e-h, Supplementary Table 4), the highest number of DEGs were found in a cluster of deep layer excitatory neurons – ExN10_L46 (238/447, 53%) and a cluster of astrocytes – Ast1 (98/447, 22%). A summary of the proportions of upregulated versus downregulated genes and unique DEGs versus DEGs shared across clusters is provided in Figure 3e. Correlations between gene expression fold differences calculated in our reanalysis and our previous analysis are provided in Supplementary Figure 8i-l (methods: Differential expression analysis - Comparison of male differential expression results to previous results).

**Figure 3:**
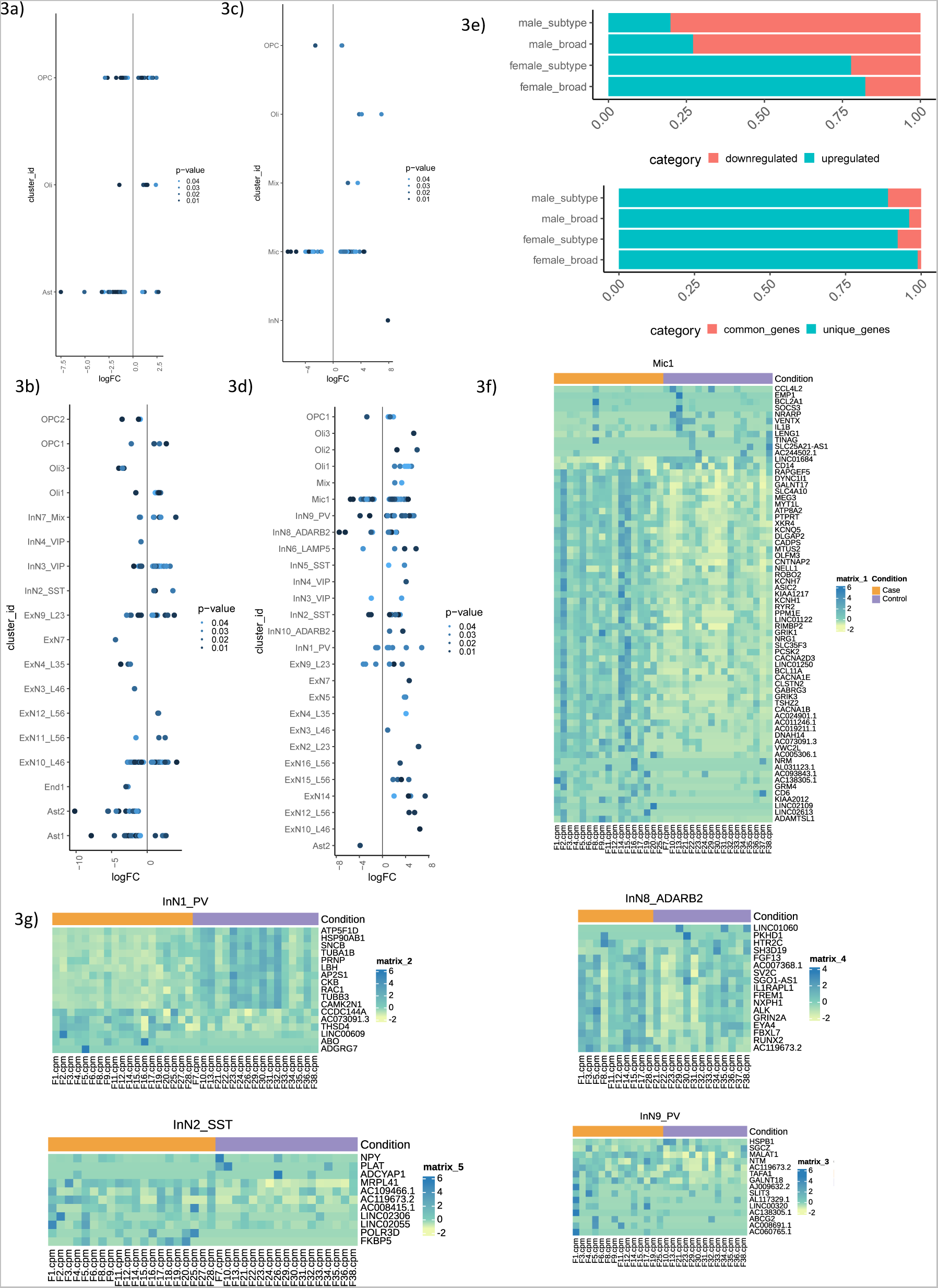
a-b) Distribution of differentially expressed genes in (a) broad cell types and (b) clusters with increased and decreased expression in male cases compared to controls. c-d) Distribution of differentially expressed genes in (c) broad cell types and (d) clusters with increased and decreased expression in female cases compared to controls. For a-d, points are colored by the corrected p-value for differential expression, and upregulated genes are plotted to the right of the midline while downregulated genes are plotted to the left. e) Barplots showing proportions of up and downregulated genes and unique and shared genes. For males, the majority of DEGs were decreased in expression in cases compared to controls both at the broad (110/151, 73%) and cluster levels (358/447, 80%) and most DEGs were cell type specific both at the broad (145/151, 96% unique DEGs) and cluster (398/447, 89% unique DEGs) level. For females, the majority of DEGs were upregulated both at the broad (70/85, 82%) and the cluster level (140/180, 78%) and most DEGs were cell type specific both at the broad (84/85, 99% unique DEGs) and cluster (166/180, 92% unique DEGs) level. f-g) Heatmaps showing the pseudobulk expression of differentially expressed genes in top clusters with highest number of DEGs in the female cluster level analysis – (f) microglia, (g) inhibitory neuronal clusters. For f-g, the plotted values are pseudobulk CPMs calculated with edgeR and muscat and scaled per row (by gene). For all heatmaps (f-g), the annotation bar at the top is colored orange for cases and purple for controls, and rows and columns are not clustered.

In females, for broad cell types, we detected a high number of DEGs in microglia only (74/85, 87%) (Figure 3c, Supplementary Figure 9a-d, Supplementary Table 5). The same analysis at the cluster level (Figure 3d, Supplementary Figure 9e-h, Supplementary Table 5) consistently showed the highest number of DEGs in the Mic1 (Figure 3f; 68/180 DEGs, 38%) cluster with a large proportion (53/68, 78%) overlapping with the microglial DEGs at the broad level. We focused on cluster level results for follow up analyses (methods: Differential expression analysis, for justification) and assessed the robustness of our microglial findings against misclassified or contaminating cells (Supplementary Figure 9i).

The majority of microglial DEGs (47/68, 69%) were confirmed to be both transcribed and translated in microglia using a TRAP gene expression dataset in a lipopolysaccharide challenge mouse model^24^.

In addition to microglia, several inhibitory neuronal clusters (Figure 3g), including two *PVALB* expressing clusters – InN1_PV and InN9_PV as well as an *SST* expressing cluster – InN2_SST and an *ADARB2* expressing cluster – InN8_ADARB2 contained the majority of remaining DEGs. Our results thus pointed to dysregulation of microglia and inhibitory neurons, especially PV interneurons in females with MDD which further prompted us to explore the biological pathways within and possible interactions between these cell types which could be altered in MDD, as detailed below.

### Meta-analysis reveals additive effects of depression-associated transcriptomic changes in males and females

To maximize statistical power to observe gene expression differences common to both males and females, we performed meta-analyses of the male and female data within each broad cell type and cluster. For broad cell types, the meta-analysis revealed upregulated genes in microglia and downregulated genes in astrocytes, with the majority of DEGs from the separate male and female analyses retained (Figure 4, Supplementary Table 6). There were more DEGs in microglia (172 DEGs) than observed in the female dataset alone (74 DEGs), whereas there were fewer DEGs in astrocytes (53 DEGs) than identified in males alone (90 DEGs). 49/90 (54%) DEGs in the broad astrocytic cluster in males and 56/74 (76%) DEGs in the broad microglial cluster in females were recapitulated in the meta-analysis. There were 22 DEGs in OPCs in the meta-analysis, but the number was less than half compared to the independent analysis of the male dataset (54 DEGs) whereas for oligodendrocytes the number of DEGs was higher when the data were meta-analyzed (21 versus 7 DEGs in the male dataset alone). The decrease in number of MDD-associated DEGs in OPCs when combining the male and female cohorts indicates that gene expression differences in OPCs in MDD are dissimilar between the sexes. This agrees with the discordance of depression-related transcriptomic changes between sexes in OPCs in our RRHO analysis.

**Figure 4:**
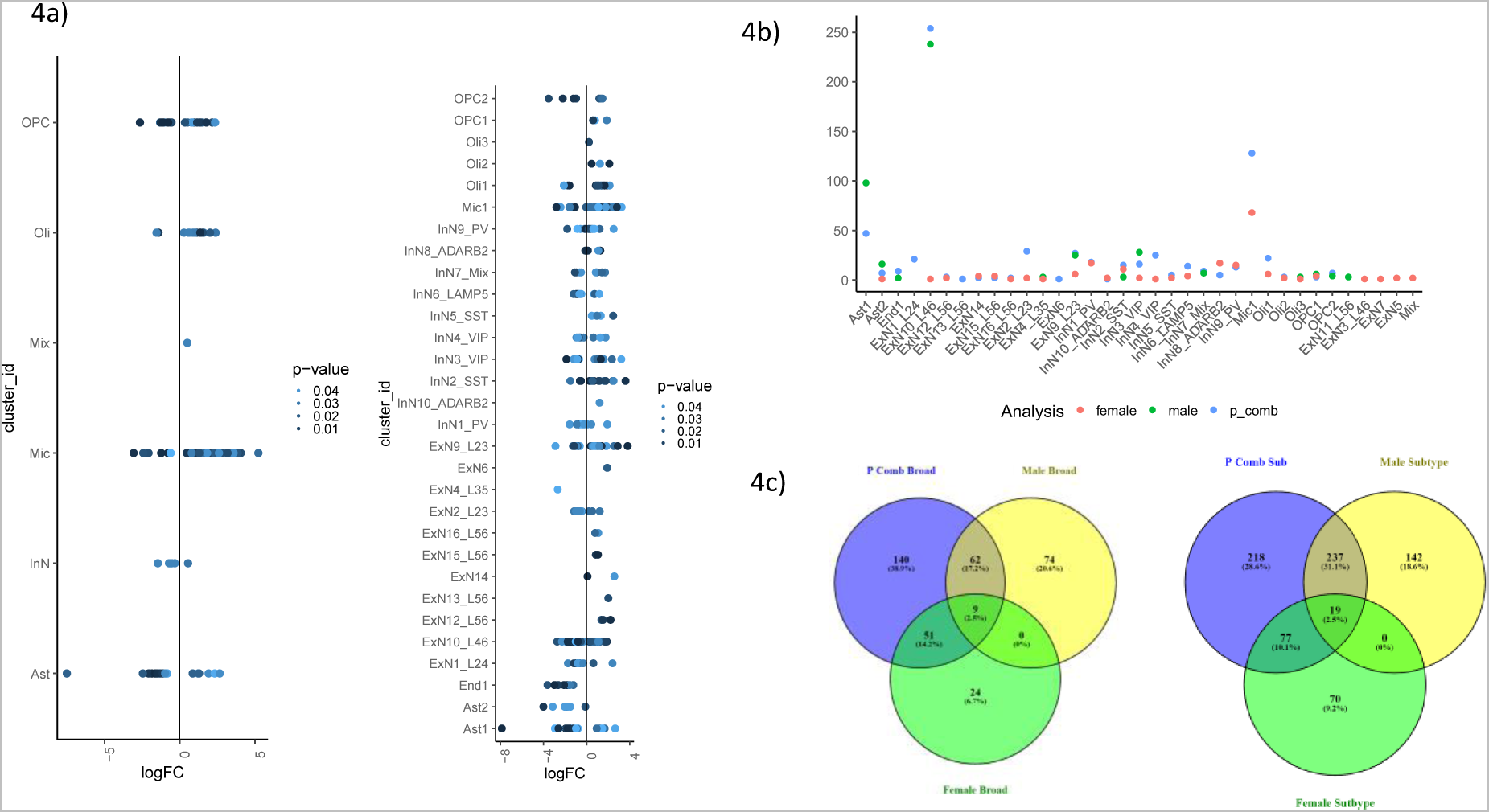
a-b) Distribution of DEGs across the (left) broad cell types and (right) clusters after p-value combination meta-analaysis. b) Numbers of DEGs in each clsuter for the male analysis, female analysis, and meta-analaysis. c-d) Overlap of meta-analysis DEGs with the individual analyses of the male and female datasets for (c) broad cell types and (d) clusters.

At the cluster level, we found that upregulated DEGs in Mic1 and downregulated DEGs in ExN10_L46 stood out as the top findings in the meta-analysis (Figure 4, Supplementary Table 6). Once again, we found more microglial DEGs (128 DEGs) via the meta-analysis compared to the female data alone (68 DEGs) and more DEGs in ExN10_L46 (254 DEGs) than with the male data alone (238 DEGs).

Given the overall between-sex concordance in MDD-associated gene expression changes detected in RRHO, it is not surprising that clusters with prominent differential expression from the individual cohorts also stood out in the meta-analysis. Taken together these results further support that the global patterns of change in gene expression within cell type are generally consistent between males and females, especially for excitatory neurons and microglia, with a few notable exceptions such as OPCs.

### Female cell type specific DEGs are enriched for previous MDD-linked genes

The relevance of the DEGs we have identified to psychiatric disorders was evaluated by referring to the PsyGeNET^25^ text-mining database. Compared to other disorders, depressive disorders had the most gene-disease associations with the female cell type specific DEGs (> 60; Figure 5a). The next largest number of gene-disease associations was for schizophrenia (< 40). Statistically, the overlap of all DEGs at the cluster level with disease-associated genes in PsyGeNET was significant only for two disease categories, Depressive disorders (hypergeometric test, p = 0.0378) and Alcohol use disorders (hypergeometric test, p = 0.0141). Further, for the top 5 clusters with highest numbers of DEGs in the female cluster-level analysis, gene-disease associations for depression and related disorders in PsyGeNET were identified for several DEGs (Figure 5b-c). Therefore, our cell type specific DEG findings in females recapitulated previously reported gene-disease associations.

**Figure 5:**
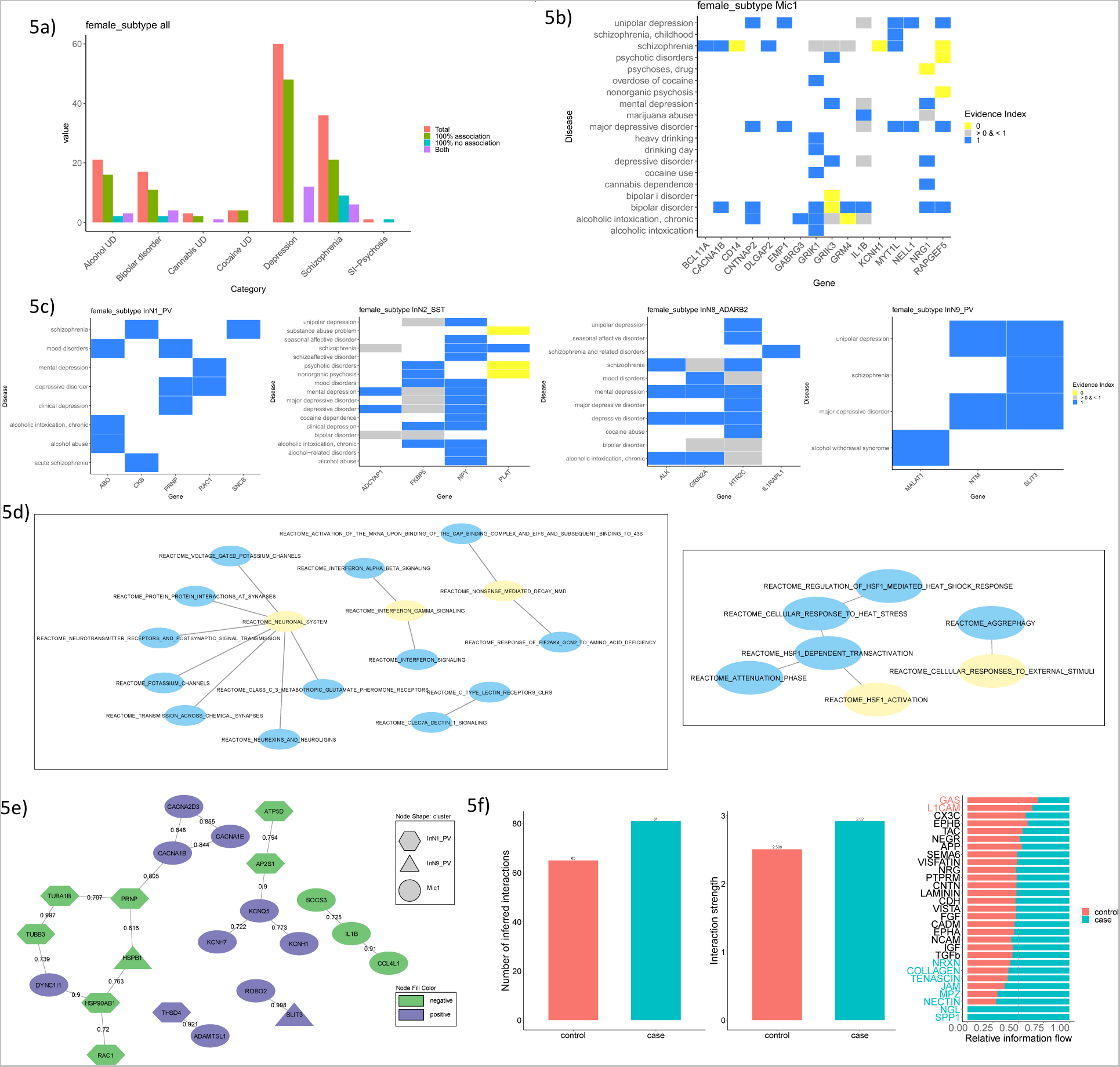
a) PsyGeNET literature reported gene-disease association bar plot for all DEGs in the female cluster level analysis. “100% association” indicates all evidence is in support, “100% no association” indicates the opposite, while “Both” indicates mixed support. b-c) Gene-disease association heatmaps for 5 clusters with the highest numbers of DEGs in females: (b) microglia, (c) inhibitory neuron clusters. Evidence index of 1 indicates that all literature supports the association, while 0 indicates that there is no support for the association. Values in between indicate partial support. d) Networks showing the relationship between main gene sets (yellow) and all gene sets (blue) with enrichment in pre-ranked GSEA with Reactome pathways in Mic1 (left) and InN9_PV (right) in females. Controlling for the overlap between gene sets, the main gene sets are independently enriched. e) STRING network showing DEGs in female microglia and PV interneurons whose protein products have reported interactions. The shape of the node represents the cluster in which the DEG was detected, and the color represents the direction of fold change in cases compared to controls. The numbers on the edges represent the confidence scores for the interactions. f) (left) Bar plots showing the number and strength of ligand-receptor communications within and between PV interneurons and microglia in cases and controls. (right) Relative strength of communication in different signaling pathways for cases and controls.

### Disease-relevant biological pathways revealed by cell type specific transcriptomic changes in females with MDD

To explore the underlying pathways associated with the cell type specific transcriptomic changes in females with MDD, we performed pre-ranked gene set enrichment analysis (GSEA; methods: Pre-ranked gene set enrichment analysis). Female microglia from cases showed significant negative enrichment scores for inflammation related Reactome pathway gene sets including “Interferon Gamma signaling”, “Interleukin 4 and Interleukin 13 signaling”, “Interleukin 10 signaling”, and “TNFR2 non-canonical NF-KB pathway” (Figure 5d, Supplementary Table 7). “Neuronal system” gene sets were positively enriched with contributions from “Voltage-gated potassium channels”, “Class C/3 metabotropic glutamate/pheromone receptors”, and “Neurexins and neuroligins” among others (Figure 5d). Interestingly both pro- and anti-inflammatory immune signaling pathway gene sets were downregulated which may indicate that MDD-associated dysregulation of gene expression in microglia involves more than just a microglial inflammatory response.

Further, both PV interneuron clusters showed a negative enrichment of heat shock factor 1 (HSF1) related terms – “HSF1 activation” in InN9_PV and “HSF1 dependent transactivation” in InN1_PV. Moreover, both clusters showed an enrichment of the gene sets “Cellular response to external stimuli” and “Metabolism of RNA”. The InN1_PV cluster showed further enrichment of immune gene sets such as “Innate immune system”, “Adaptive immune system”, and “Cytokine signaling in immune system” and interestingly in the context of sex-differences in depression, “ESR mediated signaling”, pertaining to the estrogen receptor.

Thus, our GSEA analysis of the female microglia and PV interneuron differential expression results revealed dysregulated Reactome pathway gene sets which are functionally relevant in these cell types and plausibly associated with sex-differences.

### Assessing the relationship between microglia and PV interneuron dysregulation in females with MDD

#### Protein-protein interaction assessment

To further assess the functional relevance of striking gene expression differences in microglia and PV interneurons in females with MDD, we examined whether the protein products of DEGs in these clusters belonged to interacting networks. STRING^26^ protein-protein interaction (PPI) analysis (methods: STRING analysis) revealed links between the protein products of several DEGs in the microglia and the PV interneurons. We focused on the top two interactions, based on the STRING confidence score. These interactions were between protein products of DEGs coming from microglia and PV interneurons and had the same direction of change (Figure 5e). The *ROBO2* gene, which encodes a canonical cell migration guidance receptor^27^, was increased in microglia whereas one of its corresponding ligands, *SLIT3*^27^ was increased in expression in the InN9_PV cluster. Additionally, *ADAMSTL1* and *THSD4* (also known as *ADAMTSL6*), two members of the ADAMTS-like family of proteins, which have extracellular matrix (ECM) binding properties^28^, were upregulated in microglia and in the InN1_PV cluster respectively. The PPI network analysis results point to the intriguing possibility that changes in communication between microglia and PV interneurons through contribute depression-associated brain pathology in females.

### Ligand-receptor interaction assessment

Building upon the indications from PPI assessment we explored the possible changes in ligand-receptor expression in microglia and PV interneurons between female cases and controls with CellChat^29^ (methods: CellChat analysis). CellChat identified more interacting ligand-receptor pairs and estimated increased communication strength overall within and between microglia and PV interneurons in cases compared to controls (Figure 5f). CellChat further identified several signaling pathways (groups of related ligand-receptor pairs) with decreased (top pathway: GAS) and increased (top pathway: SPP1) communication in cases compared to controls (Figure 5f). Within these top signaling pathways, we specifically identified a probable increase in SPP1 to integrin communication and decrease in GAS6-MERTK communication from microglia to PV interneurons and vice versa, respectively (Supplementary Figure 10).

### WGCNA confirms MDD dysregulated pathways in female microglia and PV interneurons

Next, we performed weighted-gene co-expression network analysis (WGCNA) using the pseudobulk gene expression profiles to identify correlated modules of genes associated with MDD in microglia and PV interneurons.

In microglia, 8 modules out of 44 had a significant correlation with case-control status (p-value < 0.05; Figure 6a). Further, the MEturquoise module which is positively correlated with MDD-status (correlation 0.627, p = 7.26×10^−5^) showed a significant overlap (p = 5×10^−56^; methods: Weighted gene co-expression network analysis) with upregulated DEGs in microglia in female cases (Figure 6b). MEturquoise also showed an enrichment of Reactome pathway gene sets related to ion channels, neurotransmitter receptors, and the neuronal system (Figure 6c) similar to gene sets found upregulated in microglia in female cases by GSEA.

**Figure 6:**
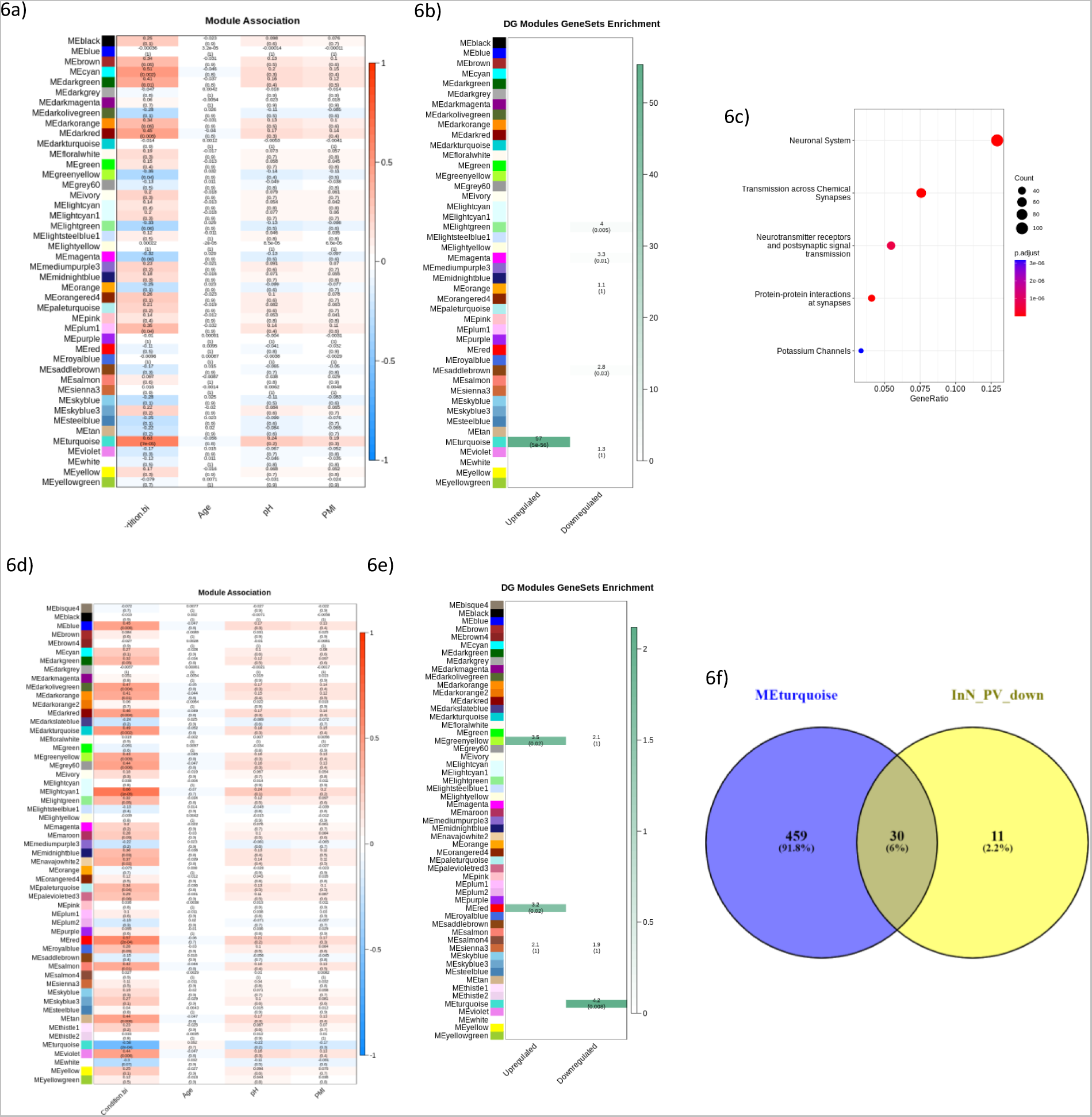
a) Heatmap showing the correlation and associated p-value, in parentheses, of Mic1 WGCNA module eigengenes with case-control status and covariates. b) Heatmap showing the test-statistic and p-value, in parentheses, for Fisher tests of overlap between the Mic1 WGCNA module member genes and DEGs in females in Mic1. c) Top Reactome pathway gene sets over-represented in Mic1 WGCNA, in the MEturqouise module. e) Heatmap showing the correlation and associated p-value, in parentheses, of InN_PV WGCNA module eigengenes with case-control status and covariates. f) Heatmap showing the test-statistic and p-value, in parentheses, for Fisher tests of overlap between the InN_PV WGCNA module member genes and DEGs in females the InN1_PV or InN9_PV clusters. g) Venn diagram showing the overlap of Reactome pathway gene sets enriched in the InN_PV WGCNA module MEturquoise (associated negatively with case status) and downregulated via GSEA in cases within InN1_PV or InN9_PV.

In PV interneurons (including nuclei in the InN1_PV and InN9_PV clusters), 16 of 55 modules were significantly associated with case-control status (p < 0.05, Figure 6d). Additionally, downregulated DEGs from InN1_PV and InN9_PV significantly overlapped (p = 0.008) with the genes from the MEturquoise module (Figure 6e). The MEturquoise module which is negatively associated with MDD (correlation −0.582, p = 0.00016), had over-representation of 489 Reactome pathway gene sets. Of these, 30 pathways overlapped with the main downregulated pathways previously identified with GSEA in InN1_PV or InN9_PV (Figure 6f). The overlapping pathways included “HSF1 activation”, “HSF1 dependent transactivation”, and “ESR mediated signaling”. Further, upregulated DEGs in InN1_PV and InN9_PV significantly overlapped with the genes of two modules which had a positive association with MDD-status: MEred (correlation 0.568, p = 0.0002) and MEgreenyellow (correlation 0.426, p = 0.0085).

Overall, the female microglia and PV interneuron WGCNA results further support our MDD-associated DEG and Reactome Pathway findings in these clusters.

## Discussion

### Sex-specificity of depression-associated transcriptomic differences

Two studies have previously evaluated sex specificity in gene expression differences associated with depression^7,10^. These studies, which analyzed bulk tissue samples, reported distinct gene expression differences in males and females with very limited overlap of DEGs. Our findings are consistent with these studies in that the cell-types with most prominent differences in gene expression – and the DEGs within these cell-types – were quite separate for males and females. However, in contrast, we found that within each cluster and broad cell type the threshold-free patterns of MDD-associated difference in gene expression were highly concordant between the sexes in most cases, except for most oligodendrocyte lineage clusters. This overall agreement between the sexes was confirmed by a meta-analysis of the male and female data. Possibly, by using single-nucleus methodology, our results provided better resolution for threshold free analyses.

Notably, recent reviews on the sex specificity of transcriptomic differences in MDD^22,30^ suggest that in females with MDD there is reduced microglial activation and increased synaptic connectivity while the opposite is true for males. This theory is supported by the downregulation we observed in MDD females in microglial inflammatory pathways such as interferon and NF-KB signaling.

### Microglial contributions to MDD in females

We found striking transcriptomic differences in microglia between females with and without MDD, while there were no differences surviving transcriptome-wide threshold in the male microglia. This salient implication of microglia specifically in females is consistent with differences in the distribution, structure, transcriptome, and proteome of microglia between the sexes, in both health and disease^31–33^. Furthermore, the number and phenotype of microglia differ by sex in the rodent brain^34–36^, and several recent rodent studies demonstrated sex-specificity of microglial response to stress in various brain regions^37–39^. These studies describe changes in genes involved in cellular stress and immune function with brain-region and sex-specific variation. This is roughly analogous to the female-specific pathway dysregulation we observed in microglia and PV interneurons in MDD.

Most studies examining peripheral markers report increased inflammation in MDD^40^, but studies in brain tissue have reported increases^40^, decreases^41^, or changes in both directions^42^ in the expression of pro-inflammatory cytokines. Moreover, several depression-linked genetic variants in pro-inflammatory genes, including in *IL1B*, *TNFA*, and *CRP*, are associated with decreased expression^43^. Recently the concept of a pro-inflammatory versus anti-inflammatory state of microglia has been challenged^44^. In the brain, amidst close interactions with multiple cell types, microglia adopt more diverse states with varying levels of pro- and anti-inflammatory markers, and this is being underscored by single-cell data^44^. Our results reflect altered microglial transcription in MDD females versus controls, with pro-inflammatory (interferon and NF-KB signaling) and anti-inflammatory (IL4, IL13, and IL10 signaling) pathways simultaneously downregulated.

We observed evidence that further “neuronal” pathways – including neurotransmitter signaling and ion channels – were upregulated in female MDD microglia, in both differential expression and WGCNA results. Microglia have long been known to express neurotransmitter receptors and ion channels. Mounting evidence suggests these canonically “neuronal” gene products regulate microglial activity^45–48^, and our results suggest that changes in their expression may contribute to MDD pathophysiology, at least in females.

### PV interneuron and microglia crosstalk in females with MDD

Together with striking changes in microglial gene expression, we observed dysregulation in PV interneurons. PV interneurons, among other interneuron subtypes, are implicated in stress and depression with evidence for sex-specific changes^49,50^. Most PV interneurons are encapsulated by ECM structures called perineuronal nets (PNNs) which help protect them from cellular stress, and microglia are known to regulate PNNs^51^. Oxidative and cellular stress relate to PV neuron and PNN deficits in animal models^52^ and cellular stress may be part of the molecular pathology in MDD^53^.

We found evidence of dysregulated cellular stress pathways, such as heat-shock factor activation, in PV interneurons in MDD females via differential expression analysis and WGCNA. Moreover, both analyses pointed to dysregulation of estrogen receptor mediated signaling. The expression of many genes is regulated by the ligand-bound estrogen receptor and difference in estrogen levels are known to contribute to differences in brain physiology between the sexes^31^.

Beyond effects in individual cell types, our results imply potentially impaired communication between PV interneurons and microglia in females with MDD. Microglial synaptic regulation involves migration of microglia towards specific neurons and in glioblastomas this migration can be regulated by SLIT-ROBO signaling^27^. The *SLIT3* gene and its corresponding receptor gene *ROBO2* were upregulated in PV interneurons and microglia respectively in females with MDD. Of note, genetic variation in *SLIT3* has been associated with depression^54^.

We observed that the ECM-binding protein genes *ADAMTSL1* and *THSD4* were upregulated in microglia and PV interneurons, respectively, in MDD females. These recently characterized ADAMTS-like proteins lack the enzymatic domains through which ADAMTSs break PNN components, but they have been proposed to protect these components from degradation by mimicking ADAMTS binding^28,55^. We therefore conjecture that microglial migration cued by PV interneurons, followed by concerted alterations of the ECM by these two cell-types stabilize PNNs in females with MDD. A recent study – including males and females – reported increased PNN number in the prefrontal cortex of MDD subjects who experienced early life adversity^56^, and our molecular findings might underlie one sex-specific mechanism for PNN alterations in MDD. Our preliminary assessment also points to downregulation of PV interneuron to microglia signaling via GAS6-MERTK and upregulation of SPP1 to integrin signaling in the opposite direction in females with MDD. Together, MERTK and GAS6 promote homeostasis and neuronal survival and they are disrupted in several nervous system disorders^57^. On the other hand, microglial osteopontin (SPP1), promotes remyelination in multiple sclerosis and is neuroprotective near infarcts in stroke but in Alzheimer’s disease it is part of the “disease associated microglia” signature^58^. The role of these signaling molecules in depression, if any, are yet to be determined.

### Limitations

This study has limitations that should be considered. We could not directly compare male and female cell type specific transcriptomes or assess the interaction of sex and disease status given that we are using data from two sex-specific datasets. Thus, the implication of different cell types in MDD between males and females could be partly attributable to differences in methodology for generating the two datasets. However, we attempted to mitigate this by applying a unified pre-processing pipeline and joint definition of cell types. Our findings are consistent with previous evidence for sex-specific mechanisms for depression etiology in animal models and human studies^6,9,59,60^.

Although our study included data from over 160,000 nuclei, the number of subjects was small relative to the large number of genes tested for associations with MDD. This may have reduced our ability to identify some disease-relevant genes and pathways.

We did not identify a separate sub-population of disease-associated microglia as observed in some neurological disorders^61^. This may partly be due to the lack of cytoplasmic transcripts in snRNA-seq limiting the information about microglial states^62^. Nevertheless, a recent study highlighted similarities between cellular and nuclear microglia RNA-seq data from mouse and human – fresh and frozen – CNS samples^63^. Nuclear microglia transcriptomes are a reliable proxy for cellular transcriptomes and are less affected by cell isolation-based transcriptional artifacts^63^. We were able to detect inflammatory pathway dysregulation in female microglia despite the limitations.

Our CellChat and STRING results are speculative. We cannot draw conclusions about the proximity of microglia to PV interneurons or the presence of PNNs, as snRNA-seq involves dissociation of the tissue with loss of spatial and structural information. Neither can we conclude that protein expression is changed for our DEGs. Future studies using spatial transcriptomic techniques coupled with immunohistochemistry may better answer these questions.

### Conclusions

We provide the first cell type and sex-specific assessment of transcriptomic changes in the dlPFC in MDD using snRNA-seq. Our dataset represents a rich resource which will stimulate further fruitful investigations of sex- and cell type specific molecular pathways in depression. While most transcriptomic changes in males with MDD are observed in deep layer excitatory neurons, astrocytes, and OPCs, in females the changes are concentrated in microglia and PV interneurons. Although major dysregulated cell types and genes are distinct for each sex, within broad cell types and clusters the patterns of transcriptomic differences in MDD are primarily concordant between males and females. Finally, preliminary evidence hints that in females with MDD, impaired communication between microglia and PV interneurons may be an important feature of MDD molecular pathology.

## Acknowledgments

GT holds a Canada Research Chair (Tier 1) and is supported by grants from the Canadian Institute of Health Research (CIHR) (FDN148374 and ENP161427 (ERA-NET ERA PerMed)). MS works in a unit funded by the UK Medical Research Council (MC_UU_00011/5).

We acknowledge the expert help of the Douglas–Bell Canada Brain Bank staff. This Project has been made possible with the financial support of Health Canada, through the Canada Brain Research Fund, an innovative partnership between the Government of Canada (through Health Canada) and Brain Canada, and in part by funding from the Canada First Research Excellence Fund, awarded to McGill University for the Healthy Brains for Healthy Lives initiative, and from the Fonds de recherche du Québec - Santé (FRQS) through the Quebec Network on Suicide, Mood Disorders and Related Disorders. The present study used the services of the Molecular and Cellular Microscopy Platform (MCMP) at the Douglas Institute.

This research was enabled in part by support provided by Calcul Québec (https://www.calculquebec.ca/en/) and the Digital Research Alliance of Canada (alliancecan.ca).

We thank Senthil Kumar Duraikannu Kailasam and the Canadian Centre for Computational Genomics team at McGill University for their services and assistance with differential expression analysis. We thank Claudia Belliveau for her suggestions on data interpretation.

## Methods

### Male snRNA-seq dataset

We used published snRNA-seq data from a cohort of male subjects with or without MDD^1^. We started with the raw FASTQ files available through GEO (<u>GSE144136</u>) and reprocessed the data, dropping two runs from one subject (number 25) with low quality results based on the previous analysis.

### Post-mortem brain samples in the female cohort

This study was approved by the Douglas Institute IRB. Human post-mortem dorsolateral pre-frontal cortex tissue was obtained from the Douglas-Bell-Canada Brain Bank (www.douglasbrainbank.ca) and from the University of Miami Miller School of Medicine Brain Endowment Bank (https://med.miami.edu/programs/brain-endowment-bank (Supplementary Table 1). Informed consent from next of kin was obtained for each individual included in this study. Frozen histological grade samples of gray and white matter were dissected from the dlPFC (Brodmann Area 9) by expert neuroanatomists and stored at –80 °C. Psychological autopsies were performed using proxy-based interviews complemented by medical charts, as previously described^2^. A summary of sample demographic characteristics is provided in Table 1. All cases included in this study died while affected by MDD, whereas controls were neurotypical individuals who died suddenly without prolonged agonal periods and did not have evidence of axis I disorders. The post-mortem interval (PMI) represents the delay between an individual’s death and collection and processing of the brain.

### Nuclei extraction, single-nuclei capture, and library preparation for female cohort

Nuclei were extracted from coronal cryosections or tissue shavings across the cortical layers and white matter, weighing between 40-65 mg, obtained using a cryostat at −20 °C with thickness set to 100 microns. Nuclei were extracted as previously described^3^. Two versions of the iodixanol gradient were used –a weaker gradient using 17.5% and 15% (w/v) concentrations of iodixanol (batches 3F, 7F, 2F) and a stronger gradient using the 29% and 25% (w/v) concentrations of iodixanol (batches 6F, 8F, 12F), as previously published^4^, and we found the stronger gradient to perform better. Nuclei were resuspended in wash buffer and stained using Hoescht 33342 (1:2000). 10 uL of nuclei were loaded onto EVE cell counting slides (MBI) and imaged using an Olympus VS120 Slide Scanner (10X magnification) and counted using the QuPath^5^ software (version 0.2.0) with the “Watershed cell detection” functionality.

We used the 10x Genomics Chromium controller for single-cell gene expression to isolate single nuclei for downstream RNA library preparation with 10x Genomics Chromium Single Cell 3’ reagents. For samples processed with version 2 of the Chromium chemistry (Supplementary Table 1), we followed the protocols as outlined by the user guide (CG00052_SingleCell3_ReagentKitv2UserGuide_RevB; latest version at https://bit.ly/3dUNOLZ), whereas for sample processed with version 3 of the Chromium chemistry (Supplementary Table 1) we followed the protocols as outlined by the user guide (CG000204_ChromiumNextGEMSingleCell3_v3.1_Rev_D, https://assets.ctfassets.net/an68im79xiti/1eX2FPdpeCgnCJtw4fj9Hx/7cb84edaa9eca04b607f9193162994de/CG000204_ChromiumNextGEMSingleCell3_v3.1_Rev_D.pdf). The only modification was for loading concentration, which we increased by 30% as we assessed the capture of nuclei to be slightly less efficient than cell encapsulation. Nuclei were loaded to capture 3000 per sample, but because of a systematic error in counting the actual number of nuclei captured per sample was variable (Supplementary Table 1).

### Sequencing, alignment, and generation of count matrices

The majority of samples in the female cohort (36) were sequenced using the Illumina NovaSeq 6000 but two samples were sequenced using BGI DNB-seq technology. Sequencing metrics are provided in Supplementary Table 1. All samples from the male cohort were realigned. Alignment was performed and count matrices were generated with Cell Ranger version 5.0.1 against the GRCh38 reference available on the 10X Genomics website (refdata-gex-GRCh38-2020-A, https://support.10xgenomics.com/single-cell-gene-expression/software/release-notes/build). We ran the “cellranger count” command using the “--include-introns” option and all other options set to default.

An initial 174,178 nuclei were obtained with Cell Ranger default cell filtering. The median value of mean reads per cell was 71,279, the average mapping rate to the transcriptome was 68.8%, the average fraction of reads in cells was 71.0 %, and the average sequencing saturation was 78.5% (Supplementary Table 1). There was higher intronic mapping rate (Kruskal-Wallis test p-value 0.0029) and a lower exonic mapping rate (Kruskal-Wallis p-value 0.0048) for cases compared to controls, but no significant differences in any other sequencing quality control metrics (Supplementary Table 1).

The filtered gene barcode matrices were individually loaded into R^6^ (versions 4.0.2 and 4.1.2) for downstream analysis and processed with Seurat^7^ (4.0.3.9000 and 4.0.5). Percentage of reads from mitochondrially encoded genes were calculated before filtering, added as metadata, and used as a quality control parameter for nuclei filtering, after which the mitochondrial genes were removed for downstream analysis. The parameters for filtering were as follows:

Male cohort: nCount_RNA < 35000, nFeature_RNA > 350, percent.mt < 10

Female cohort v2 chemistry: nCount_RNA < 25000, nFeature_RNA > 250, percent.mt < 10

Female cohort v3 chemistry: nCount_RNA < 120000, nFeature_RNA > 350, percent.mt < 10

After filtering, we obtained 79,058 nuclei in the male cohort (43,347 from cases, 35,711 from controls) and 81,653 nuclei in the female cohort (49,926 from cases, 31,727 from controls). In the female cohort, after filtering, the median across samples of the median number of UMIs per cell and the median the number of genes per cell were 2758.5 and 1711.5 respectively (Supplementary Table 1). In the males, the corresponding numbers were 2530.5 and 1638.25 respectively (Supplementary Table 1).

### Dimensionality reduction and data integration

We performed SCTransform on each Seurat object individually and used the SelectIntegrationFeatures function to set the variable genes for downstream analysis. We scaled each cell to 10000 counts and ran log normalization. We regressed out nCount_RNA and percent.mt from the counts to get scaled gene expression values for variable genes, which was used as input for calculating 100 PCA components. We corrected PCA components with Harmony^8^ to account for batch, chemistry, and sample specific effects. This helped align the datasets as seen in the UMAP projections produced before and after correction (Supplementary Figure 1a-d). All UMAPs in figures were created using Seurat.

### Clustering

We tested of a range of combinations of clustering parameters for the Seurat package (FindClusters function) using the scclusteval^9^ sub-sampling (80% of all cells, 100 times) and stability comparison workflow using Jaccard indices. With each sub-sampling, PCA and Harmony were recalculated. The parameters tested were: k-param: 20, 30; resolution: 0.1, 0.3, 0.5, 0.7, 0.9, 1.1, 1.3, 1.5; number of Harmony corrected PCs to use: 70, 80.

We then set a threshold for the minimum stability with a chooseR-like^10^ approach based on the bootstrapped medians of the median Jaccard index across all the clusters and all the parameter sets tested. We selected parameters that maximized the number of clusters while passing the threshold of cluster stability: 70 Harmony corrected PCA components, a k-nearest neighbors’ parameter of 30, and a resolution of 0.7 (Supplementary Figure 2a-b). Repeating the Harmony correction with a seed set followed by clustering with the optimal parameters produced 41 clusters. Final UMAPs were produced using all 100 Harmony corrected PCA components and all calculation parameters set to default.

### Cluster annotation

Genes enriched in clusters were calculated using the wilcoxauc function from presto^11^ with default parameters, and filtered with the following criteria: padj < 0.05, logFC > log(1.5), pct_in-pct_out > 10. For annotation, the following known cell type marker genes were assessed in the cluster enriched genes:

Macrophage/microglia: *SPI1, MRC1, TMEM119, CX3CR1;* Endothelial: *CLDN5, VTN, VIM;* Astrocytes: *GLUL, SOX9, AQP4, GJA1, NDRG2, GFAP, ALDH1A1, ALDH1L1;* OPCs: *PDGFRA, PCDH15, OLIG2, OLIG1;* Oligodendrocytes: *PLP1, MAG, MOG, MOBP, MBP;* Neurons: *SNAP25, RBFOX3;* Excitatory neurons: *SATB2, SLC17A7, SLC17A6;* Inhibitory neurons: *GAD1, GAD2, SLC32A1,* Inhibitory neuronal subtypes: *VIP, PVALB, SST, ADARB2, LHX6, LAMP5, PAX6*

Additionally, expression of cell-type specific genes from BRETIGEA^12^ were assessed using the Seurat AddModuleScore function (Supplementary Figure 4).

Twenty clusters of excitatory cells were identified (Supplementary Figure 5a) including four superficial cortical layer neuronal clusters (ExN1_L24, ExN2_L23, ExN8_L24, ExN9_L23), ten deep cortical layer neuronal clusters (ExN3_L46, ExN4_L35, ExN10_L46, ExN11_L56, ExN12_L56, ExN13_L56, ExN15_L56, ExN16_L56, ExN19_L56, ExN20_L56) and six excitatory neuronal clusters without an obvious pattern of cortical layer specific marker expression (ExN5, ExN6, ExN7, ExN14, ExN17, ExN18). The layer annotations of excitatory neuronal clusters were supported by assessment of enrichment for genes known to be specific to the different layers of the cortex using spatial transcriptomics results form Maynard et al.^13^ (data in supplementary table 4 of the cited publication).

We identified 10 inhibitory clusters (Supplementary Table 2, Supplementary Figure 5b), that can broadly be divided into cells likely derived from the medial ganglionic eminence (MGE; InN1_PV, InN9_PV, InN2_SST, InN5_SST) based on *LHX6, SST,* or *PVALB* enrichment, or the caudal ganglionic eminence (CGE; InN3_VIP, InN4_VIP, InN6_LAMP5, InN8_ADARB2, InN10_ADARB2) based on *ADARB2* enrichment. The InN2_SST cluster was enriched for *SST* and *GAD1* expression but had no *LHX6* enrichment. The InN8_ADARB2 cluster also showed enrichment for *SST*. One inhibitory neuron cluster with enrichment for both *ADARB2* and *LHX6* (InN7_Mix), which has been previously reported^14^.

### Assessment of clustering quality

Contribution of batches, groups, and subjects was relatively uniform across clusters (Supplementary Figure 3a-d). Endothelial, microglial, and oligodendrocyte lineage cells showed a higher percentage of contribution from the females compared to the males, possibly due a different dissection strategy used for the two cohorts such that for the female cohort more white matter tissue was included in the nuclei extractions. All but one cluster (number 34, later annotated at ExN17) had contributions from both the male and female cohorts and one cluster was primarily composed of cells from the female cohort and showed exceptionally high numbers of UMIs detected per nucleus (number 11, ExN5) (Supplementary Figure 3f). Moreover, one cluster (number 17, which was later annotated as showing a mixed expression profile – Mix) had relatively high percentage of mitochondrial reads (Supplementary Figure 3f).

### Comparison to other datasets

#### MetaNeighbor

We used MetaNeighbor^15^ to compare the clusters in our dataset to several published datasets^1,16,17^. For the Song et al., 2020 data we used the h5_a^18^, h7^19^, h10^20^, and h14^14^ datasets which contain adult human cortical cells or nuclei and were reprocessed by the authors. We used our own dataset as a reference to train the model, for consistency of comparisons across the datasets and limited the analysis to the same variable genes we used for PCA and clustering. MetaNeighbor best hits plots are shown in Figure 1e and Supplementary Figure 6. Using the published datasets as the reference showed similar correspondence between the clusters (data not shown).

#### Spatial label transfer

We used Seurat to transfer the labels for layer annotation from a spatial transcriptomics dataset^13^ to our dataset (Supplementary Figure 5a). Each tissue section of the spatial transcriptomics data was treated separately and one section each from two different subjects were assessed (data shown for one subject and section - 151673). Both the spatial and snRNA-seq data were preprocessed with SCTransform and transfer anchors were identified using the “canonical correlation analysis” option before transferring the labels.

### Pseudotime trajectory analysis

We used “slingshot”^21^ to build a pseudotime trajectory with our OL nuclei (Supplementary Figure 5c). We built the pseudotime trajectory with the male and female datasets combined. OL nuclei were subset and UMAP was rerun using the following parameters: dims= 1:10, min.dist=0.1, spread = 5, n.neighbors = 100, chosen to capture the global patterns in the data. Slingshot was run with the resulting UMAP as input and using the following parameters: extend = “n”, start.clus = “OPC2”, end.clus = “Oli3”, stretch = 0.1, thresh = 0.3, once again chosen to capture the broad patterns in the data. The start and end clusters were chosen based on their position in the UMAP, and cluster labels were provided. The oligodendrocyte lineage (OL) clusters were arranged from OPC2 at one end of the pseudotime trajectory, followed by OPC1 and OPC3, a small cluster possibly corresponding to committed oligodendrocyte precursors (COPs). At the other end of the pseudotime trajectory Oli2, Oli1, and Oli3 were placed sequentially and could represent the order of oligodendrocyte clusters from myelinating to mature states.

We fit the expression of genes along pseudotime by splitting the data for males and females before using tradeSeq^22^ (Supplementary Figure 5d). We ran fitGAM on the UMI counts for each gene, with age, PMI, pH, and batch as covariates, with conditions set to case and control status, and nknots of 5, based on evaluation of a range of nknot values. The fitted expression of OL marker genes was visualized using the plotSmoothers function. The pseudotime trajectory analysis was performed following the vignette available here: https://kstreet13.github.io/bioc2020trajectories/articles/workshopTrajectories.html

### Cell type proportions comparison

The percentage of nuclei in each cluster and each broad cell type for each sample was calculated and compared between cases and controls with Wilcoxon tests using rstatix^23^. Further to mitigate the effect of outliers we obtained p-values for the Wilcoxon test using bootstrapping with 10000 replicates (R package boot^24^; Supplementary Table 3) which supported the initial results. Lastly, we also examined the distribution of p-values (Supplementary Table 3) from the Wilcoxon test rerun after randomly sub-sampling 70% of the nuclei 100 times similar to a previous study^25^ and confirmed the pattern of changes in proportion preserved after sub-sampling (Supplementary Table 3).

### Differential expression analysis

We performed pseudobulk differential gene expression analysis using muscat^26^ and edgeR^27^ at the broad cell type and cluster levels in males and females separately. Pseudobulk expression profiles were obtained by summing the raw UMI counts for each gene for each sample within the broad cell type or cluster. Only one run of sample 24 was included in these analyses (M24_2 excluded). Additionally, subjects were only included if they had a minimum of 10 cells in the broad cell type or 5 cells minimum in the cluster. The covariates included age, pH, PMI, and batch and muscat’s default gene and sample filtering were disabled. DEGs were selected using an FDR adjusted local (within cluster or broad cell type) p-value <0.05 and logFC > log2(1.1) and non-zero expression value in at least 3 samples. The isOutlier function (nmads 5, log “FALSE”) from scater^28^ was used to flag potential outliers on the CPMs from edgeR, as an additional assessment for genes that were called as differentially expressed. Flagged outliers were not removed from analysis.

Since the only difference between the broad and cluster level microglial results lies in the exclusion criteria for subjects based on number of cells contributed the input data and outcomes were similar between these analyses, and we focused on the cluster level results in females for follow-up analyses.

All Venn diagrams to show overlap of differentially expressed genes were made with Venny (version 2.1, https://bioinfogp.cnb.csic.es/tools/venny/index.html).

#### Comparison of male differential expression results to previous results

For the male differential expression results, using linear regression we compared the log fold changes per gene for top clusters with highest numbers of DEGs from the current analysis with the per gene estimates for similar clusters with high numbers of DEGs in our previous analysis^1^. Considering only the top 1000 genes in common ranked by the p-values in the current analysis, we found moderate positive relationships with R-squared values in the 0.13-0.32 range (Supplementary Figure 8i-l). Considering that the analysis approaches were quite distinct at every upstream and downstream step, these results support a similar pattern of changes in gene expression in the male data as we had previously reported.

#### Sub-clustering of microglia for differential expression analysis in females

A subset of microglia clustered next to oligodendrocytes in the UMAP, which could reflect misclassified cells, doublets, or even immune oligodendroglia^29,30^ or white matter microglia^31^. To determine the robustness of our microglial results to the presence of this subset of cells, we sub-clustered the microglial cluster. We found variable features within the microglial population, reran PCA and Harmony, and optimized clustering parameters (resolution 0.01, other parameters default) using silhouette scores. We excluded any subclusters which expressed oligodendrocyte lineage markers (*PLP1* and *ZFPM2*). Then we reran differential expression analysis on female microglia using the same parameters as initially used and found that new per gene logFCs showed a strong positive association (linear regression) with the initial results (Supplementary Figure 9i). Given that subsetting microglia to most confident nuclei did not substantially alter the results and for downstream analysis, we proceeded with the DEGs obtained using the full microglial cluster.

### Comparison of male and female results

#### Rank-rank hypergeometric overlap

We performed a threshold free, rank-rank hypergeometric overlap (RRHO) analysis with RRHO2^32^. Within each cluster or broad cell type, genes were scored using the product of the logFC and the negative log10 uncorrected p-value from differential expression analysis in the male and female datasets separately. The scored gene lists were provided to RRHO2_initialize function (method “hyper” and log10.ind “TRUE”) and the results were plotted using the RRHO2_heatmap function.

#### Meta-analysis by Fisher combination of p-values

To meta-analyze the male and female differential expression results per board cell type and per cluster, we used Fisher combination of p-values as implemented in the metaRNASeq^33^ R package on the uncorrected p-values after filtering out genes detected in less than 3 samples. We also used an FDR adjusted p-value threshold of 0.05 for genes to be considered significantly changed in the meta-analysis and removed any genes with opposite direction of change between the two datasets.

### Functional interpretation of female differential expression results

#### Pre-ranked gene set enrichment analysis (GSEA)

For the microglial (Mic1) and PV interneuron (InN1_PV, InN9_PV) differential expression results we individually performed pre-ranked Gene Set Enrichment Analysis^34^ with FGSEA^35^ using the same ranking metric as used for RRHO (product of log fold change and the negative log base 10 of the uncorrected p-value). We evaluated the Reactome pathway^36^ gene sets obtained from msigdbr^37^. The following parameters were used for the fgsea function: eps = 0.0, minSize = 15, maxSize = 1000 and any pathways with adjusted p-value < 0.05 were considered to be significant. Finally, we ran collapsedPathways with pval.threshold= 0.01 to get the main pathways for each cluster.

#### PsyGeNET analysis

With the list of DEGs from the female dataset across all clusters, we ran enrichedPD from psygenet2r^38^ with database = “ALL” and other parameters set to default to find the psychiatric disorders for which our DEGs showed an enrichment. Next, we ran psygenetGene with database= “ALL” and other parameters set to default, and created a geneAttrPlot for the evidence index for all DEGs from all clusters in females to summarize the links between our DEGs and psychiatric disorders reported in PsyGeNET^39^. Additionally, we similarly ran psygenetGene, individually on the DEGs from Mic1, InN1_PV, InN2_SST, InN9_PV, InN8_ADARB2, and plotted the corresponding gene-disease association heatmaps with plot type = “GDA heatmap”.

#### STRING analysis

We used STRING DB^40^ (version 11.5) to assess the relationships between the protein products of our DEGs in female microglia and PV interneurons. The entire list of DEGS from these clusters (Mic1, InN1_PV, and InN9_PV) were provided as input and the confidence level was set to high (interaction score > 0.7). We then exported the network to Cytoscape (3.9.1), colored genes by direction of change in expression, shaped DEG nodes based on their cluster of origin, and labelled the edges with the confidence scores for the interactions.

#### CellChat analysis

We subsetted the relevant nuclei from females in Mic1, InN1_PV, and InN9_PV and performed CellChat^41^ analysis. We relabelled all PV interneuron nuclei as InN_PV. For cases and controls independently, ran sequentially ran identifyOverExpressedGenes and identifyOverExpressedInteractions with lenient default parameters to find the ligand-receptor gene combinations overexpressed in these cell types. Next, we ran computeCommunProb (with nboot = 1000) followed by computeCommunProbPathway, netAnalysis_computeCentrality, and aggrgateNet with default parameters to find the ligand-receptor pathways present. Lastly, we merged the case and control objects and ran computeNetSimilarityPairwise with type “functional”. Finally, we used the compareInteractions, rankNet, and netVisual_bubble to visualize the results. We used the following vignette for CellChat analysis: https://github.com/sqjin/CellChat/blob/master/tutorial/Comparison_analysis_of_multiple_datasets.html

### Weighted gene co-expression network analysis (WGCNA)

Weighted gene co-expression network analysis (WGCNA) was performed to identify co-expression modules using the snRNA-seq expression data^42^. First, the aggregated expression for each female sample in microglia and PV interneuron clusters (InN1_PV and InN9_PV combined) was calculated by summing the counts per gene across all nuclei. We excluded subjects that did not have at least 5 microglial nuclei or 5 PV interneuron nuclei (InN1_PV and InN9_PV combined). To account for known external sample traits, the counts were corrected for age, pH, PMI, and batch (same as covariates used for differential gene expression analysis) using limma^43^. In addition, lowly expressed genes with total counts of below 5 were removed. A soft thresholding power of 10 and 12, respectively, with a minimum module size of 30 genes, were used for network construction and module detection for microglia and PV interneurons. Each module was correlated with the phenotype (healthy control vs MDD), and significance was determined using a p-value < 0.05.

To further characterize modules correlated with MDD, Fisher tests for overlap were performed to calculate the over-representation of DEGs as described previously^44^. In addition, the functional annotation of modules was determined using Reactome Pathway gene set over-representation analysis provided by clusterprofiler^45^.

