## Supplementary Figures for "Cell type specific transcriptomic differences in depression show similar patterns between males and females but implicate distinct cell types and genes"

### Supplementary Information

### Supplementary Figures

1a) 1b)

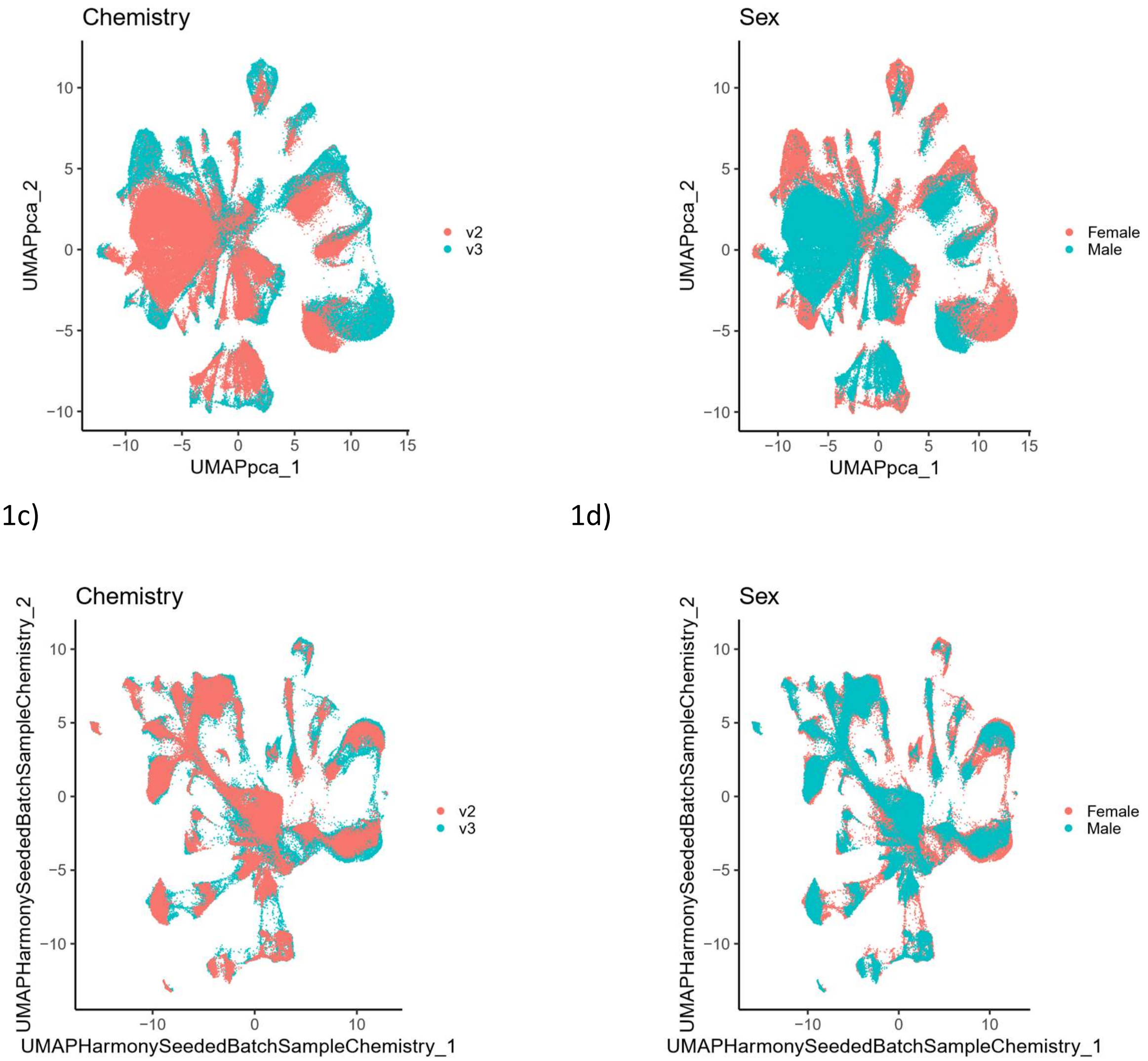


**Supplementary Figure 1:** a-b) UMAP plot using uncorrected PCA components colored by chemistry and sex. c-d) UMAP plot using Harmony corrected PCA components colored by chemistry and sex. For UMAPs we used all 100 PC components or Harmony corrected PC components with other parameters set to default.

2a)

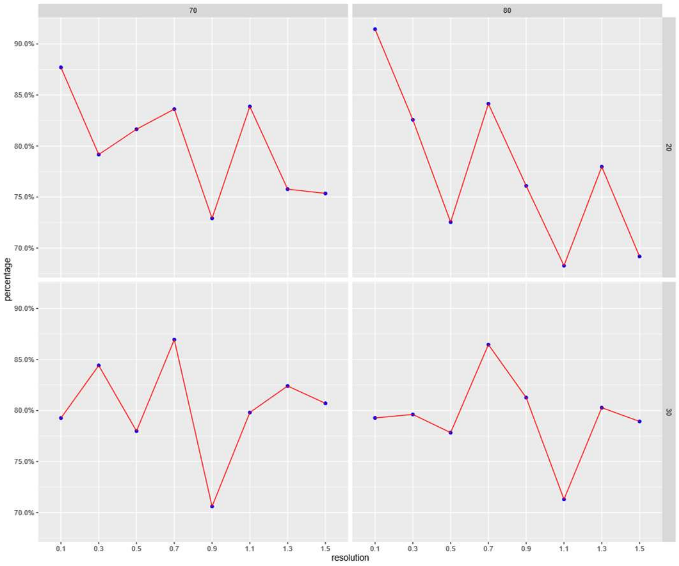


2b)

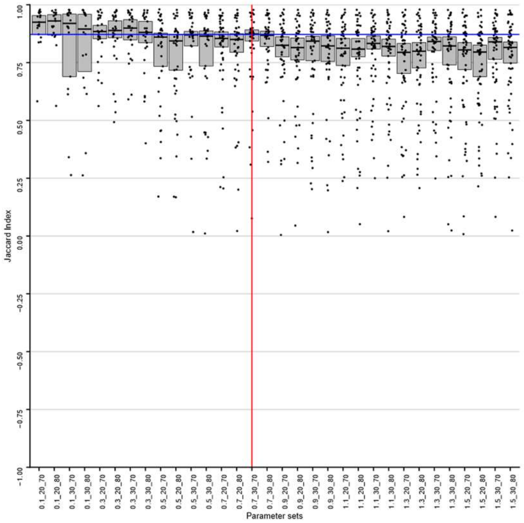


**Supplementary Figure 2:** a) scclusteval output showing the percentage of nuclei in stable clusters, as assessed by sub-sampling and Jaccard index calculation, using a range of clustering parameters. b) Boxplots showing the median Jaccard index for each cluster across sub-sampling with different parameter combinations for clustering. The clustering parameters that provide the highest number of clusters, while crossing a bootstrapped threshold of median Jaccard index across clusters, are highlighted.

3a)

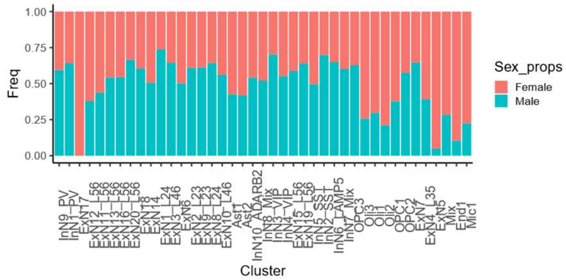


3b)

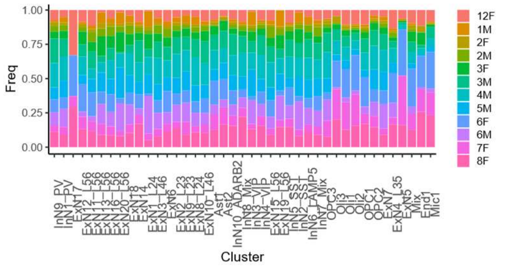


3c)

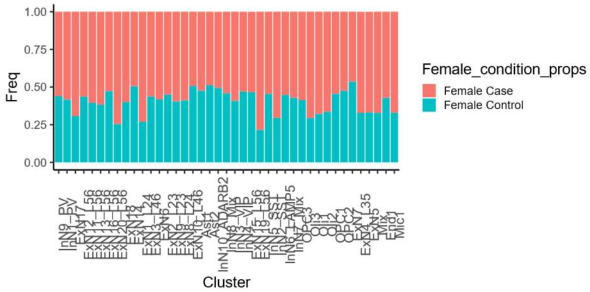


3d)

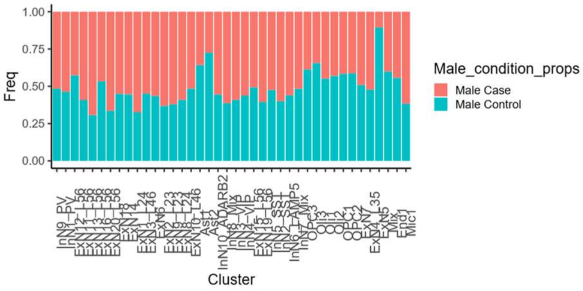


3e)

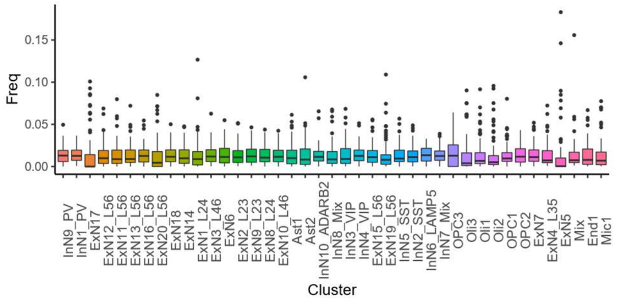


3f)

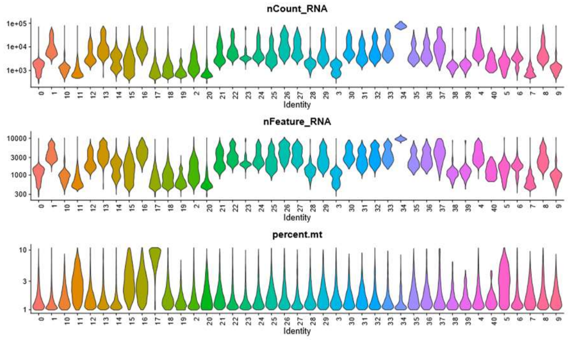


**Supplementary Figure 3:** a) Proportions of nuclei from each sex in each cluster. b) Proportion of nuclei from each batch in each cluster. c) Proportion of nuclei from cases and controls among the female nuclei in each cluster. d) Proportion of nuclei from cases and controls among the male nuclei in each cluster. e) Proportion of nuclei from each library (mostly corresponding to subject) in each cluster. f) Violin plots for number of molecules detected, number of genes detected, and percentage of mitochondrial reads per nuclei split by cluster. Mitochondrial gene counts were removed for downstream analysis, after calculating the mitochondrial read percentage. Y-axis is in log-scale. Correspondence between numbered clusters and cluster names in 3f: 0 - ExN1_L24, 1 - ExN2_L23, 2 - Ast1, 3 - Oli1, 4 - ExN3_L46, 5 - ExN4_L35, 6 - Oli2, 7 - Oli3, 8 - InN1_PV, 9 - InN2_SST, 10 - InN3_VIP, 11 - ExN5, 12 - InN4_VIP, 13 - ExN6, 14 - OPC1, 15 - End1, 16 - ExN7, 17 - Mix, 18 - Mic1, 19 - OPC2, 20 - Ast2, 21 - InN5_SST, 22 - ExN8_L24, 23 - ExN9_L23, 24 - ExN10_L46, 25 - InN6_LAMP5, 26 - ExN11_L56, 27 - ExN12_L56, 28 - ExN13_L56, 29 - InN7_Mix, 30 - ExN14, 31 - InN8_ADARB2, 32 - ExN15_L56, 33 - ExN16_L56, 34 - ExN17, 35 - InN9_PV, 36 - InN10_ADARB2, 37 - ExN18, 38 - ExN19_L56, 39 - ExN20_L56, 40 - OPC3.

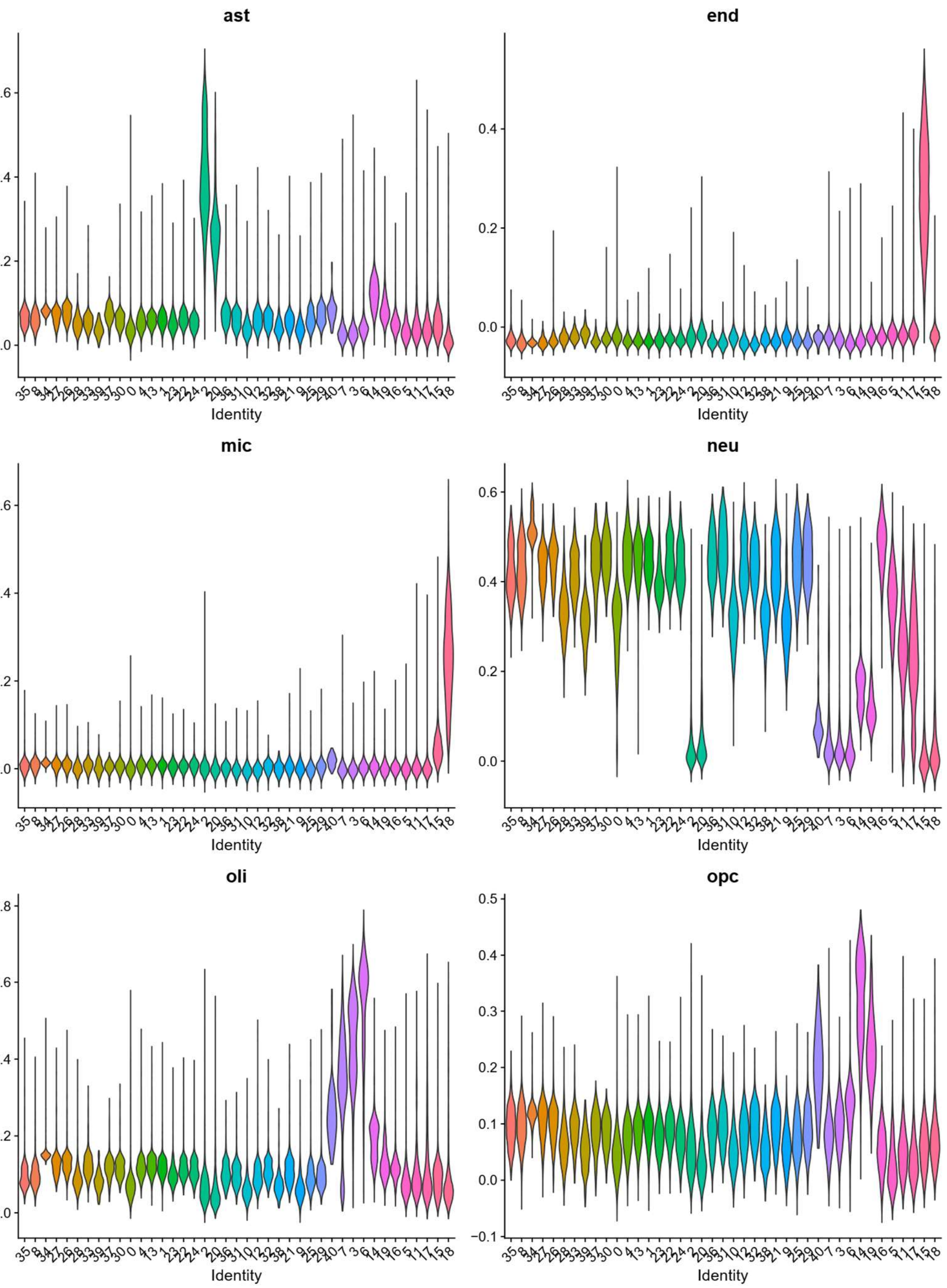


**Supplementary Figure 4**

Violin plots showing module scores for major brain cell type marker genes from BRETIGEA in each cluster. Correspondence between numbered clusters and cluster names same as in Supplementary Figure 3.

5a)

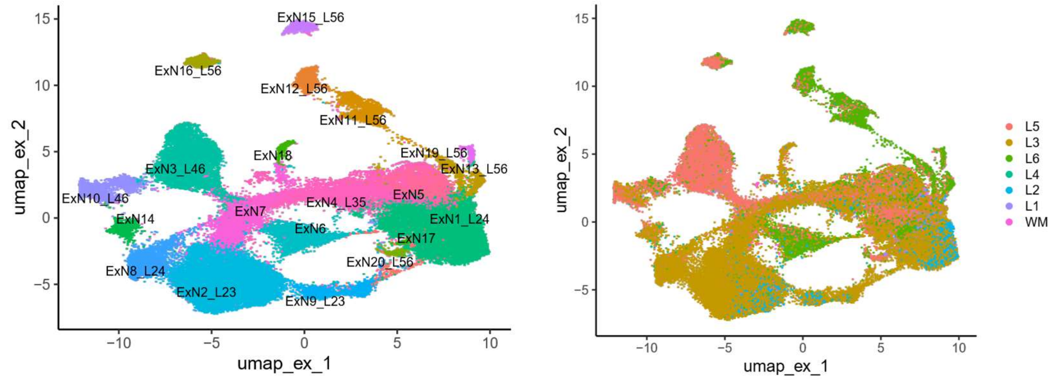
ExN15_L56
ExN16_L56
ExN12_L56
ExN11_L56
ExN19_L56
ExN13_L56
ExN18
ExN3_L46
ExN10_L46
ExN14
ExN7
ExN4_L35
ExN5
ExN1_L24
ExN6
ExN17
ExN20_L56
ExN8_L24
ExN2_L23
ExN9_L23
umap_ex_1
umap_ex_2
L5
L3
L6
L4
L2
L1
WM

5b)

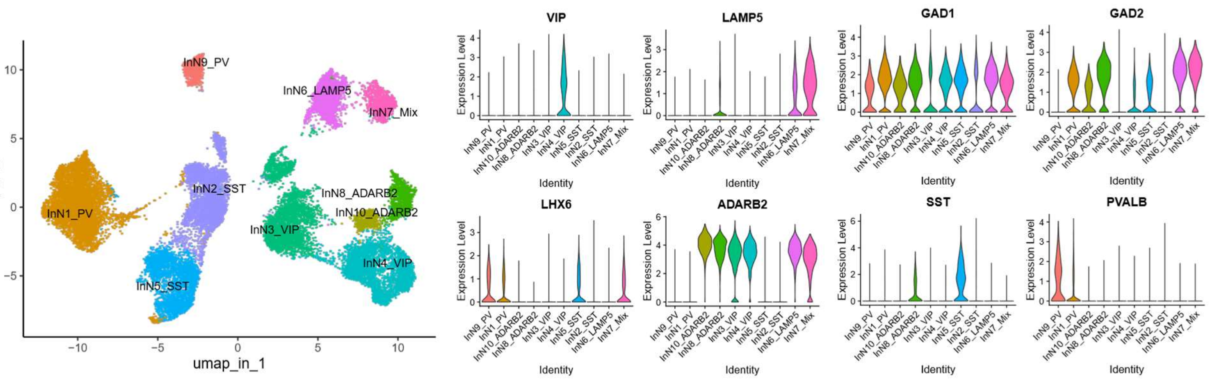
InN9_PV
InN6_LAMP5
InN7_Mix
InN2_SST
InN8_ADARB2
InN1_PV
InN10_ADARB2
InN3_VIP
InN4_VIP
InN5_SST
umap_in_1
umap_in_2
VIP
LAMP5
GAD1
GAD2
LHX6
ADARB2
SST
PVALB
Expression Level
Identity

5c)

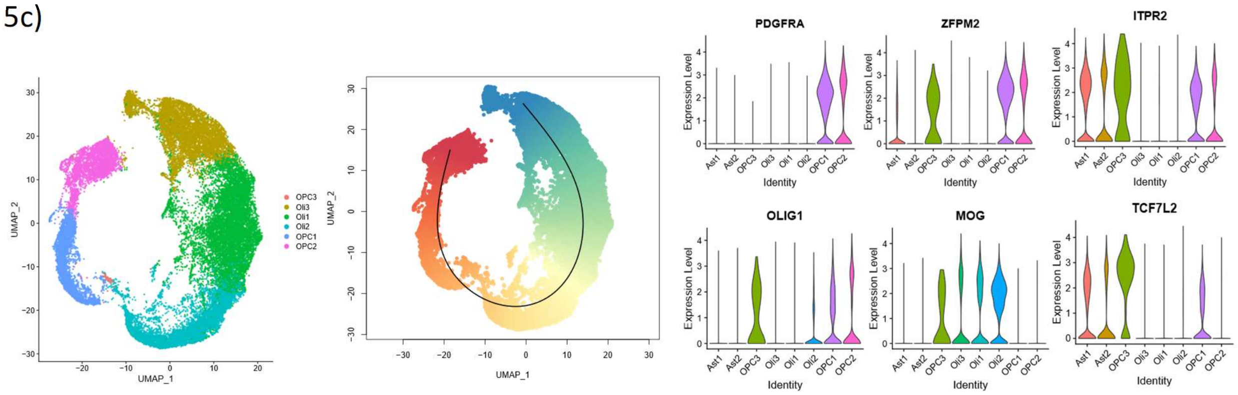
OPC3
Oli3
Oli1
Oli2
OPC1
OPC2
UMAP_1
UMAP_2
PDGFRA
ZFPM2
ITPR2
OLIG1
MOG
TCF7L2
Ast1
Ast2
Expression Level
Identity

5d)

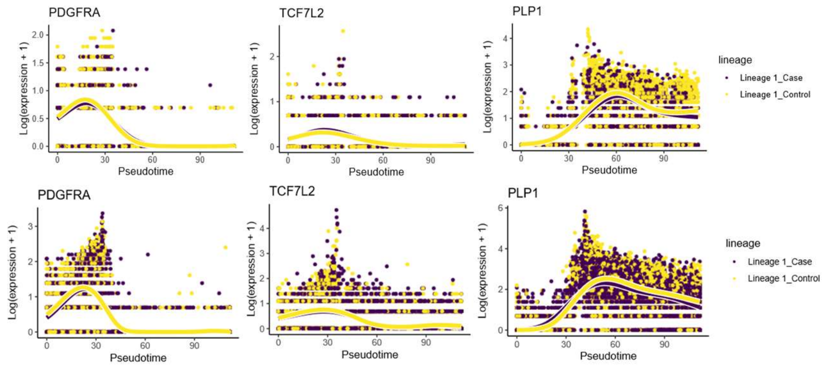
PDGFRA
TCF7L2
PLP1
Log(expression + 1)
Pseudotime
lineage
Lineage 1_Case
Lineage 1_Control

**Supplementary Figure 5.** a) (left) UMAP plot showing the 20 clusters of excitatory neurons identified. (right) UMAP plot showing the predicted layer labels for excitatory neurons using Seurat label transfer from a spatial transcriptomics dataset of the human dlPFC[1]. b) (left) UMAP plot showing the 10 clusters of inhibitory neurons identified. (right) Violin plots showing the expression of maker genes of inhibitory neurons and their known subtypes. c) (left) UMAP plot showing the 6 oligodendrocyte lineage (OL) clusters. (middle) Same UMAP plot as colored according to the pseudotime trajectory calculated (early to late pseudotime points colored from red to blue). (right) Violin plots showing expression of selected markers of the OL in the respective clusters. d) Smoothed expression fit to depict the variation in expression of selected OL genes along pseudotime using GAMS. For UMAPs in (a) and (b) we used the first 50 Harmony components, and n.neighbors = 20.

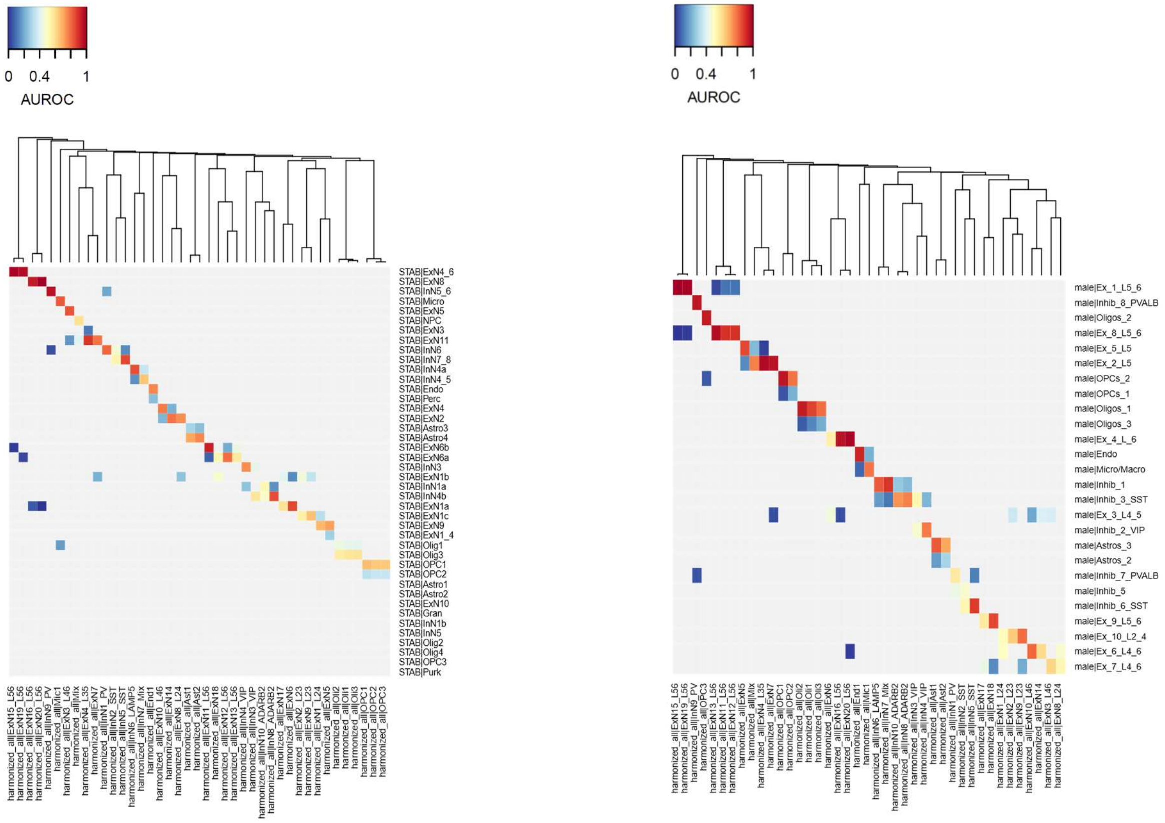


**Supplementary Figure 6**

Best hits plot from MetaNeighbor showing the correspondence between clusters defined in our dataset and clusters defined in (left) the STAB dataset and (right) in our previous analysis of the male cohort alone.

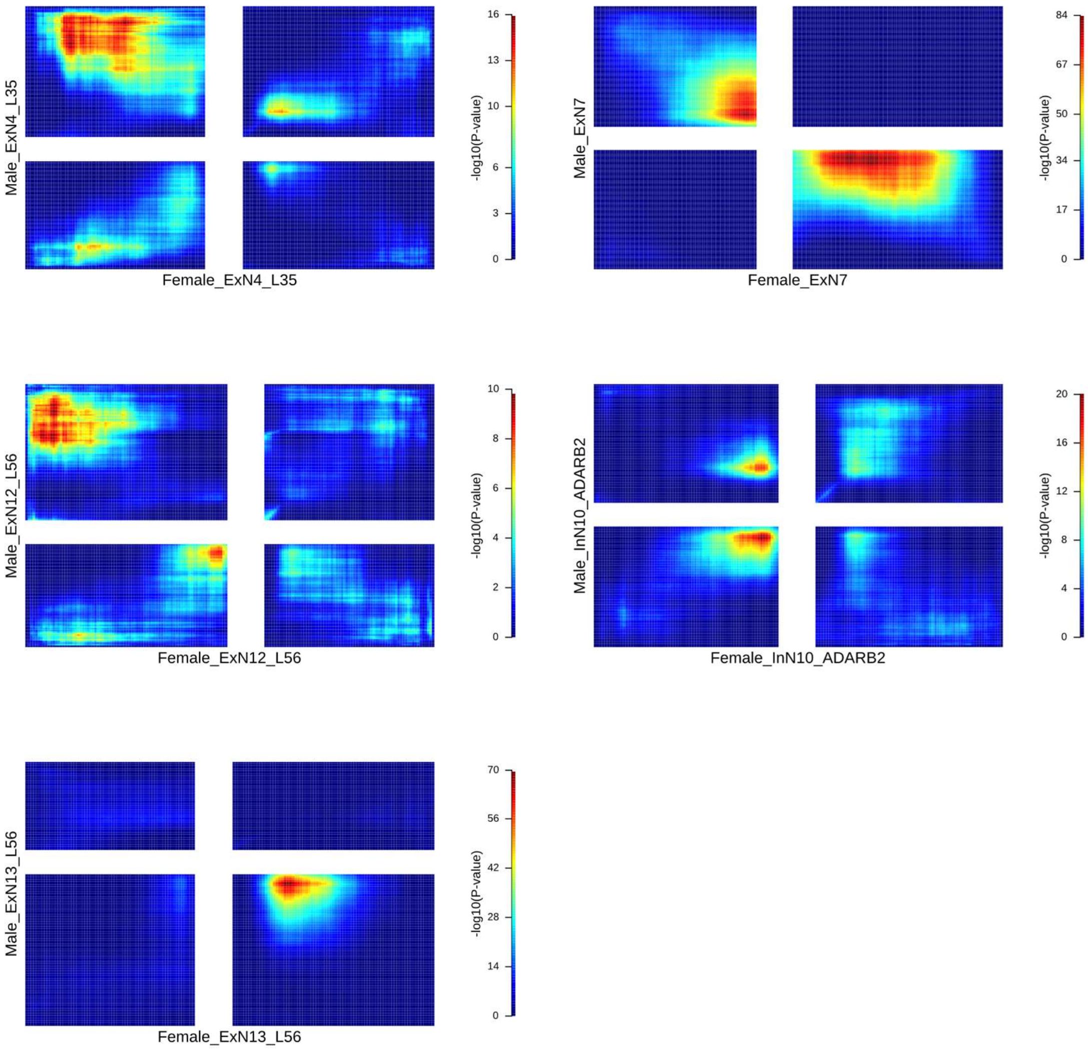


**Supplementary Figure 7**

RRHO plots showing the discordant relationship between males and females for patterns in depression-associated gene expression difference in several excitatory and inhibitory neuronal clusters.

8a)

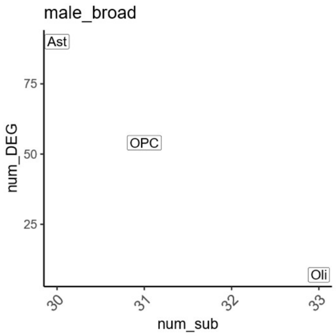


8b)

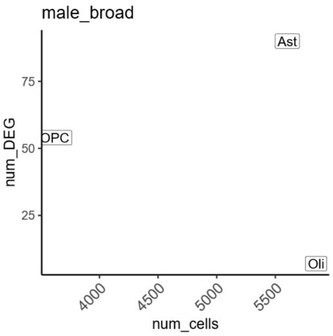


8c)

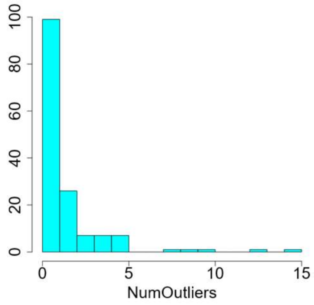


8d)

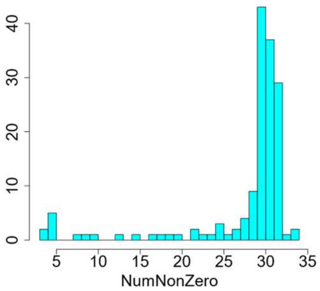


8e)

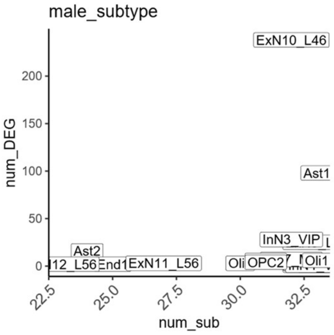


8f)

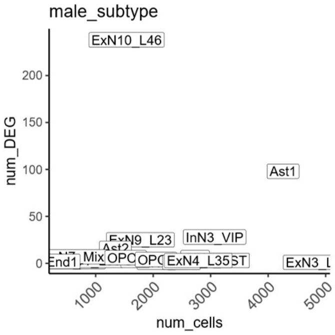


8g)

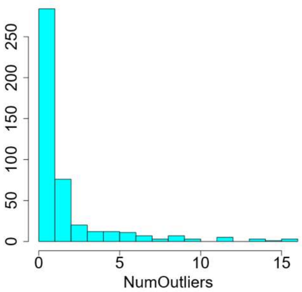


8h)

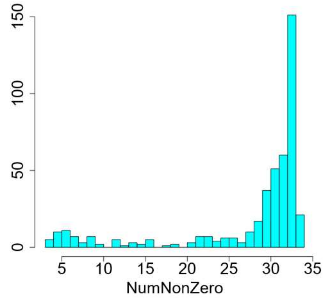


8i)

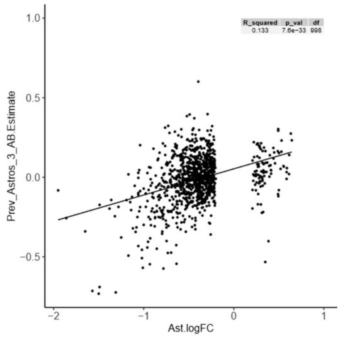


8j)

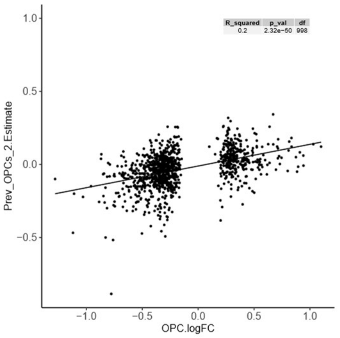


8k)

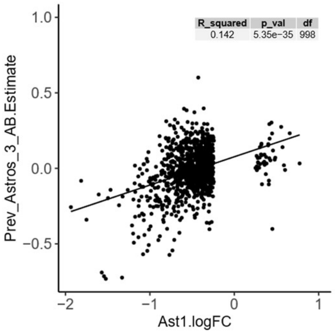


8l)

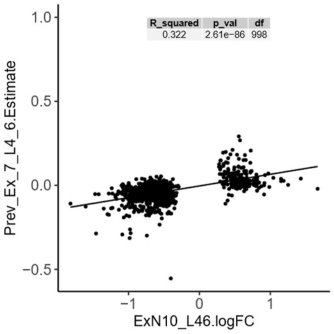

**Supplementary Figure 8:** Number of DEGs in each broad cell type for males plotted against (a) the number of subjects included in DEG analysis and (b) the total number of nuclei in the broad cell type. For DEGs in males at the broad cell type level (c) the distribution of number of subjects flagged as possible outliers and (d) the distribution of subjects with non-zero expression. Number of DEGs in each cluster for males plotted against (e) the number of subjects included in DEG analysis and (f) the total number of nuclei in the cluster. For DEGs in males at the cluster level (g) the distribution of number of subjects flagged as possible outliers and (h) the distribution of subjects with non-zero expression. Possible outliers were not removed from the analyses but were assessed as a quality metric for DEG analysis. The plots include the Mix cluster results. i-l) Scatter plots showing the relationship between (linear regression and corresponding statistics) the estimated effects per gene from our previous analysis of the male data and the log fold changes from our current analysis for cell type specific case-control differences in males for similar pairs of clusters.

9a)

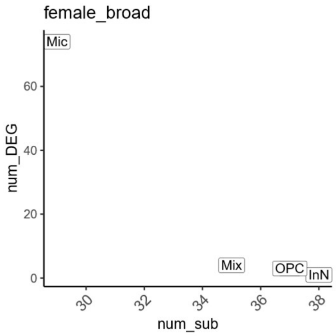


9b)

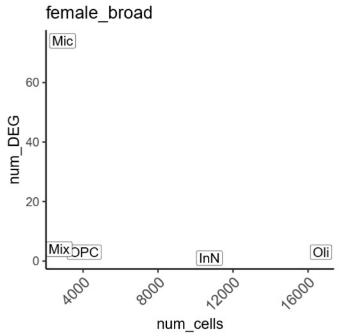


9c)

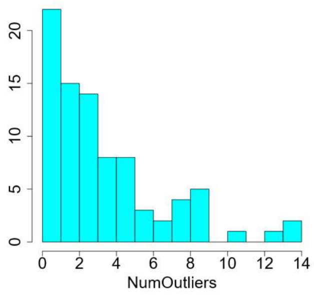


9d)

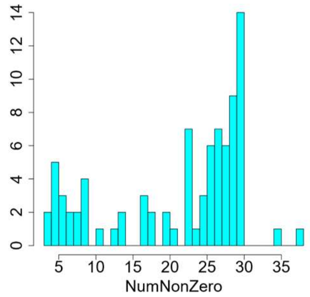


9e)

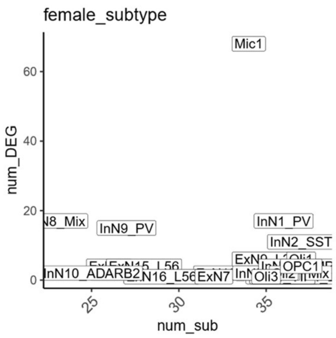


9f)

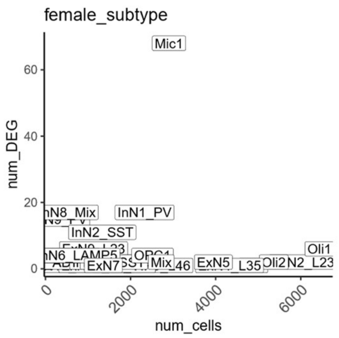


9g)

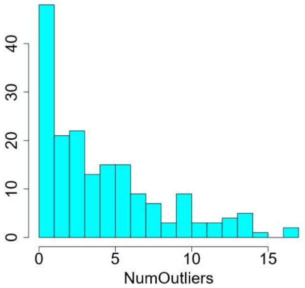


9h)

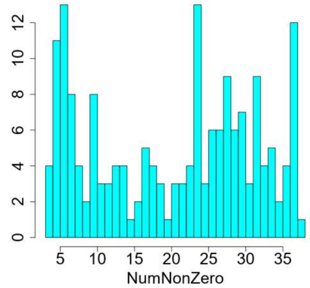


9i)

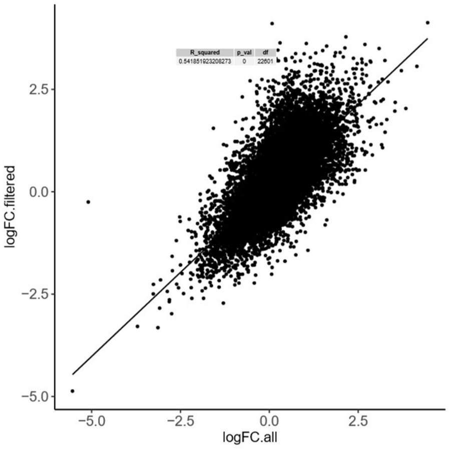

**Supplementary Figure 9:** Number of DEGs in each broad cell type for females plotted against (a) the number of subjects included in DEG analysis and (b) the total number of nuclei in the broad cell type. For DEGs in females at the broad cell type level (c) the distribution of number of subjects flagged as possible outliers and (d) the distribution of subjects with non-zero expression. Number of DEGs in each cluster for females plotted against (e) the number of subjects included in DEG analysis and (f) the total number of nuclei in the cluster. For DEGs in females at the cluster level (g) the distribution of number of subjects flagged as possible outliers and (h) the distribution of subjects with non-zero expression. Possible outliers were not removed from the analyses but were assessed as a quality metric for DEG analysis. The plots include the Mix cluster results. i) Scatter plot of log fold changes per gene in DEG analysis of the female microglia data at the cluster level with or without including nuclei that express oligodendroglial markers and cluster close to oligodendroglia on the UMAP plot (methods: Differential expression analysis - Sub-clustering of microglia for differential expression analysis in females). The majority of DEGs obtained with the full microglia cluster were retained in the subsetted microglia cluster (40/68, 59%) and the log fold changes for DEGs were strongly positively related (linear regression) with an R-squared of 0.54.

10a)

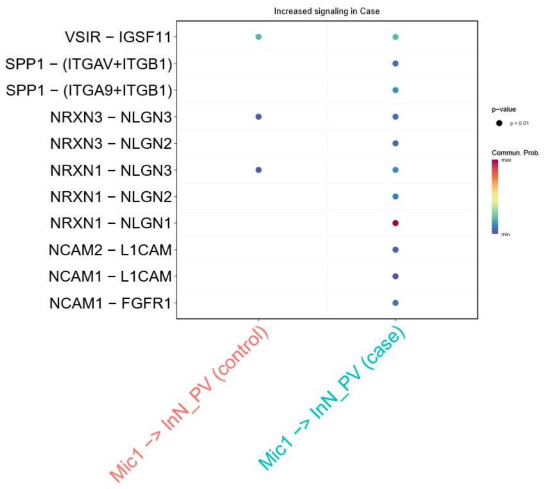


10b)

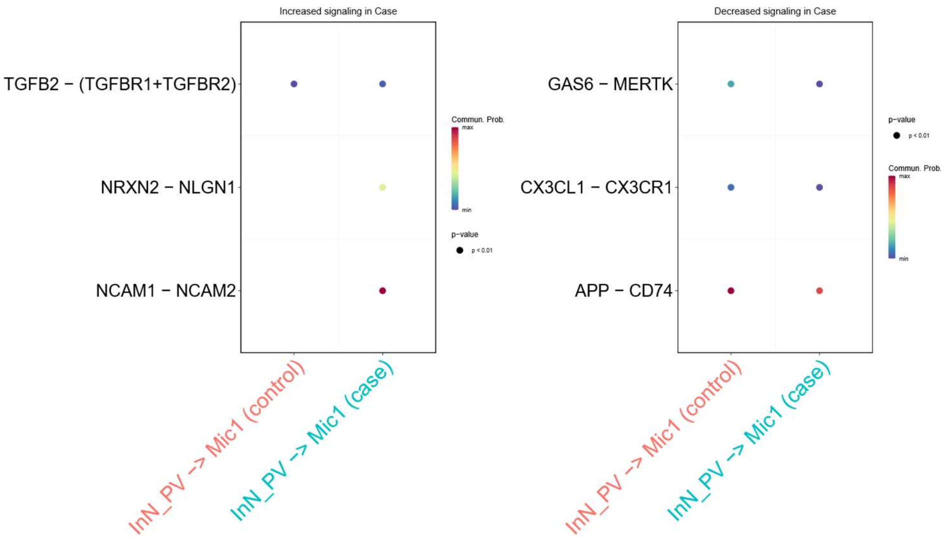


**Supplementary Figure 10:** a) Ligand-receptor pairs with increased signaling from microglia to PV interneurons in cases compared to controls. b) Ligand receptor pairs with increased (left) and decreased (right) signaling from PV interneurons to microglia in cases compared to controls. These results are based on a preliminary assessment using CellChat.
